# Paradoxical activation of oncogenic signaling as a cancer treatment strategy

**DOI:** 10.1101/2023.02.06.527335

**Authors:** Matheus Henrique Dias, Anoek Friskes, Siying Wang, Joao M. Fernandes Neto, Frank van Gemert, Soufiane Mourragui, Hendrik J. Kuiken, Sara Mainardi, Daniel Alvarez-Villanueva, Cor Lieftink, Ben Morris, Anna Dekker, Emma van Dijk, Chrysa Papagianni, Marcelo S. da Silva, Robin Jansen, Antonio Mulero-Sánchez, Elke Malzer, August Vidal, Cristina Santos, Ramón Salazar, Rosangela A. M. Wailemann, Thompson E. P. Torres, Giulia De Conti, Jonne A. Raaijmakers, Petur Snaebjornsson, Shengxian Yuan, Wenxin Qin, John S. Kovach, Hugo A. Armelin, Hein te Riele, Alexander van Oudenaarden, Haojie Jin, Roderick L. Beijersbergen, Alberto Villanueva, Rene H. Medema, Rene Bernards

**Affiliations:** Division of Molecular Carcinogenesis, Oncode Institute, The Netherlands Cancer Institute, Amsterdam, The Netherlands; Division of Cell Biology, Oncode Institute, The Netherlands Cancer Institute, Plesmanlaan 121, 1066 Amsterdam, the Netherlands; State Key Laboratory of Oncogenes and Related Genes, Shanghai Cancer Institute, Renji Hospital, Shanghai Jiao Tong University School of Medicine, Shanghai, China; Division of Tumor Biology & Immunology, The Netherlands Cancer Institute, Plesmanlaan 121, 1066 Amsterdam, the Netherlands; Hubrecht Institute-KNAW (Royal Netherlands Academy of Arts and Sciences) and University Medical Center, Utrecht, Netherlands; Division of Molecular Carcinogenesis, The Netherlands Cancer Institute, Amsterdam, The Netherlands; Chemoresistance and Predictive Factors Group, Program Against Cancer Therapeutic Resistance (ProCURE), Catalan Institute of Oncology (ICO), Oncobell Program, Bellvitge Biomedical Research Institute (IDIBELL), L’Hospitalet del Llobregat, Barcelona, Spain; Division of Molecular Carcinogenesis, NKI Robotic and Screening Center, The Netherlands Cancer Institute, Amsterdam, The Netherlands; Department of Biochemistry, Institute of Chemistry, University of São Paulo, São Paulo, SP, Brazil; Department of Chemical and Biological Sciences, Institute of Biosciences, São Paulo State University (UNESP), Botucatu, SP, Brazil; Department of Pathology; University Hospital of Bellvitge, Bellvitge Biomedical Research Institute (IDIBELL), L’Hospitalet de Llobregat, Barcelona, Spain; Xenopat S.L., Parc Cientific de Barcelona (PCB), Barcelona, Spain; Department of Medical Oncology, Catalan Institute of Oncology (ICO), Oncobell Program, Bellvitge Biomedical Research Institute (IDIBELL), CIBERONC, Barcelona, Spain; Center of Toxins, Immune-response and Cell Signaling, Instituto Butantan, São Paulo, SP, 05503-900, Brazil; Department of Clinical and Experimental Oncology, Federal University of São Paulo (UNIFESP), São Paulo, Brazil; Department of Pathology, The Netherlands Cancer Institute, Amsterdam, the Netherlands; University of Iceland, Faculty of Medicine, Reykjavik, Iceland; The Third Department of Hepatic Surgery, Eastern Hepatobiliary Surgery Hospital, Shanghai, China; Lixte Biotechnology Holdings, Inc., 680 East Colorado Blvd., Suite 180 Pasadena, CA 91101

## Abstract

Cancer homeostasis depends on a balance between activated oncogenic pathways driving tumorigenesis and engagement of stress-response programs that counteract the inherent toxicity of such aberrant signaling. While inhibition of oncogenic signaling pathways has been explored extensively, there is increasing evidence that overactivation of the same pathways can also disrupt cancer homeostasis and cause lethality. We show here that inhibition of Protein Phosphatase 2A (PP2A) hyperactivates multiple oncogenic pathways and engages stress responses in colon cancer cells. Genetic and compound screens identify combined inhibition of PP2A and WEE1 as synergistic in multiple cancer models by collapsing DNA replication and triggering premature mitosis followed by cell death. This combination also suppressed the growth of patient-derived tumors *in vivo*. Remarkably, acquired resistance to this drug combination suppressed the ability of colon cancer cells to form tumors *in vivo*. Our data suggest that paradoxical activation of oncogenic signaling can result in tumor suppressive resistance.

## INTRODUCTION

Aberrant oncogenic signaling resulting from genetic and non-genetic alterations underlies the pathological proliferation and other hallmarks of cancer^1^. The last decades brought a myriad of targeted drugs to inhibit oncogenic signaling, resulting in meaningful progress in the treatment of cancer. Unfortunately, long-lasting control of advanced cancers with these agents remains virtually elusive due to the emergence of resistance^2^. The pervasive heterogeneity and plasticity of advanced cancers result in the rapid selection of drug-resistant variants that have rewired cellular signaling such that the therapy becomes ineffective^3^. Frequently, resistance to targeted therapies results from secondary mutations that re-activate the signaling pathway in the presence of the drug. This suggests that effective control of cancer may require approaches that are fundamentally different from inhibition of oncogenic signaling.

There is increasing evidence demonstrating that hyperactivation of oncogenic signaling can be as lethal to cancer cells as the inhibition of these oncogenic pathways^4–6^. A particularly compelling example in this respect is the observation that while expression of either a mutant *EGFR* or a mutant *KRAS* oncogene in the lung epithelial cells of a mouse causes cancer, their co-expression in the lung epithelium is toxic, rather than tumorigenic^7^. Moreover, it is evident that oncogenic signaling in cancer cells is accompanied by an increased mobilization of stress response pathways to survive the stress associated with the oncogenic activity^8^. This scenario suggests that deliberate hyperactivation of oncogenic signaling pathways in cancer cells may lead to an extreme reliance on stress response pathways, creating potential vulnerabilities. We have recently reviewed the rationale for a “paradoxical intervention” in cancer treatment and discussed how this approach can potentially address current challenges in cancer therapy^9^. Such an approach consists of hyperactivation of oncogenic signaling combined with the targeting of specific stress response pathways.

While there is a vast arsenal of drugs to inhibit oncogenic signaling, the options for hyperactivation of oncogenic signaling are more limited. Protein phosphatase 2A (PP2A) is a serine/threonine phosphatase that acts in multiple cancer-relevant pathways, including mitogenic signaling, DNA damage response, and apoptosis^10–12^. Evidence that PP2A acts as a tumor suppressor in certain contexts by restraining oncogenic signaling has sparked the generation of Small Molecule Activators of PP2A (SMAPs) to reactivate the enzyme^13^. As activation of PP2A inhibits oncogenic signaling, its inhibition should further activate these pathways. LB-100, a small molecule inhibitor of PP2A has shown toxicity in various cancer models. Activation of mitogenic signaling and engagement of stress response pathways have been associated with these anti-cancer effects^14, 15^. Importantly, in a phase 1 clinical trial, LB-100 showed a rather favorable toxicity profile in doses associated with clinical response^16^. This set of characteristics makes LB-100 an attractive drug to test the concept of paradoxical activation of oncogenic signaling for the treatment of cancer.

Using colorectal cancer cells as primary models, we show here that LB-100 hyperactivates multiple oncogenic signaling pathways and simultaneously engages several stress response pathways. Concomitant inhibition of the WEE1 kinase and PP2A is highly lethal in multiple cancer models and suppresses tumor growth *in vivo*. Most strikingly, we found that acquired resistance to this drug combination was associated with loss of the tumorigenic phenotype.

## RESULTS

### LB-100 activates mitogenic signaling and engages stress response pathways in colorectal cancer cells

We initially focused on colorectal cancer (CRC) models to test the notion that the PP2A inhibitor LB-100 further activates oncogenic signaling. We selected a panel of seven CRC cell lines carrying diverse oncogenic mutations frequently found in patients (*KRAS*, *BRAF*, *APC*, *TP53*, *CTNNB1* and others)^17^ (Supplementary table S1). Dose-response assays showed moderate toxicity of LB-100 in the low micromolar range in these CRC models, with IC50s varying from 1.5 to 7.2 µM (Figure S1A).

Although we aimed to use LB-100 to activate mitogenic signaling and engage stress-response pathways, PP2A phosphatases have multiple targets^12^ and their inhibition likely has broader effects on cellular processes. To gain a comprehensive insight into the cellular processes modulated by LB-100, we treated HT-29 and SW-480 CRC cells with a sublethal concentration of LB-100 (4 µM) for 1, 4, 8, 12, or 24h and performed RNA sequencing assays. We used Gene Set Enrichment Analyses (GSEA) to compare these treated samples to their respective untreated controls. We focused the analyses on the "hallmark" and "KEGG" molecular signatures to cover a wide range of biological states or processes. Figure 1A (left) shows that LB-100 induces a transient positive enrichment in K-Ras, MAPK, mTORC1, and WNT gene sets in both CRC models, as well as the KEGG colorectal cancer gene set. These data corroborate that LB-100 further activates oncogenic pathways in CRC cells. Positive enrichment of gene sets associated with NF-κB signaling, UV response (DNA damage), Unfolded Protein Response (UPR), hypoxia, and apoptosis are also seen (Figure 1A, right), indicating the engagement of different stress response pathways in response to LB-100. Figure S1B shows the remaining hallmark and KEGG gene sets found significantly enriched in at least one time point in both cell lines. Overall, gene sets related to mitogenic signaling, stress pathways and inflammatory response pathways were positively enriched upon LB-100 treatment.

**Figure 1:**
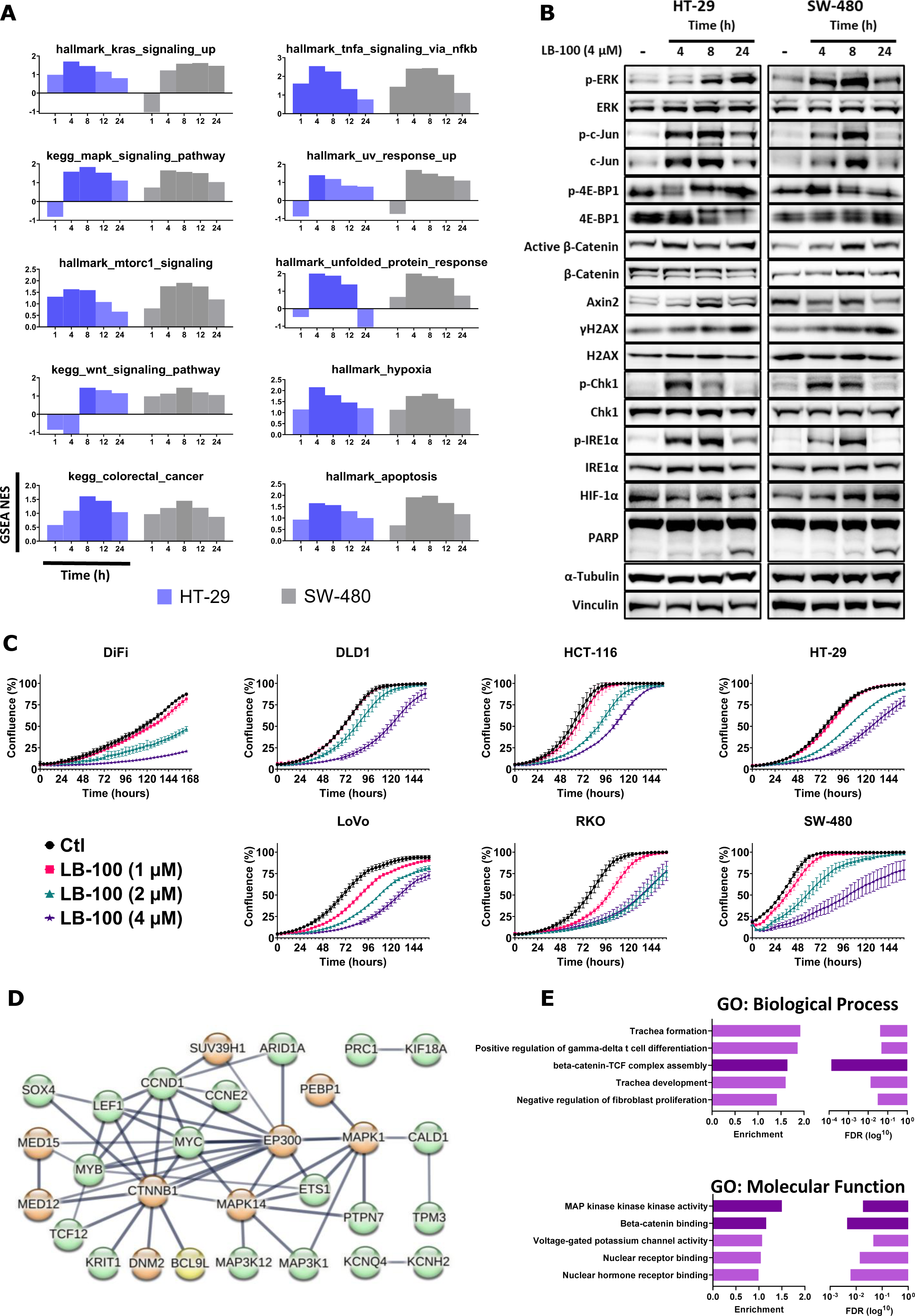
LB-100 activates oncogenic signaling, engages stress response pathways, and restrains the proliferation of CRC cells. (A) Gene Set Enrichment Analyses on time-course transcriptome data from HT-29 and SW-480 cells show selected “Hallmarks” and “KEGG” molecular signatures modulated by LB-100 (4 µM). Darker bars indicate time points for which the respective gene set were significantly enriched (p-value <0.05). **(B)** Time-course western blots show selected oncogenic signaling and stress response pathways modulated by LB-100 (4 µM) in HT-29 and SW-480 cells. α-Tubulin and Vinculin were used as loading controls. **(C)** IncuCyte-based proliferation assays with the CRC models in the absence or presence of LB-100 at 1, 2, or 4 µM for the indicated times. **(D)** String network combining all hits identified by the two independent genome-wide CRISPR screens (Figure S2) as modulators of LB-100 toxicity. Only high-confidence interactions are shown and disconnected nodes are omitted. Green nodes: CRISPRa screen; orange nodes: CRISPR-KO screen; yellow node: identified on both screens. **(E)** Gene Ontology (GO) analyses using the full list of hits from both CRISPR screens (Figure S2) as input. The top 5 enriched Biological Processes and Molecular Functions terms are shown. Darker bars highlight WNT/β-Catenin- and MAPK-related terms.

We performed a series of Western blots to validate the findings of the transcriptome analyses. Figure 1B shows that LB-100 treatment further activated the MAPK pathway as evidenced by increases in p-ERK and p-c-JUN in both cell lines. A mobility shift of 4E-BP1 (indicative of activation by phosphorylation) was noticeable after LB-100 treatment, corroborating mTORC1 activation. Moreover, AXIN2, a transcriptional target of the WNT/β-catenin pathway, was upregulated by LB-100 in HT-29 cells. Similarly, the levels of active β-catenin increased in SW-480 cells, supporting the positive enrichment of the WNT signaling gene set observed in these cell lines. The engagement of stress response pathways was also apparent at the protein level as γ-H2AX and p-CHK1, common markers of DNA damage or replication stress, increased after LB-100 treatment in both cell lines. A sharp increase in p-IRE1α, a major proxy for Unfolded Protein Response (UPR) activation, was also induced by LB-100 in these models. A noticeable increase in the master transcriptional regulator of hypoxia response, HIF-1α, was found in SW-480 cells, but not in HT-29. Furthermore, modest PARP cleavage was evident on both CRC models 24h after LB-100 treatment (Figure 1B). Despite the evident activation of oncogenic signaling pathways induced by LB-100, IncuCyte-based cell proliferation assays showed no increase in cell proliferation induced by 1, 2, or 4 µM of LB-100 in the CRC panel. Instead, impaired cell proliferation is evident across the panel (Figure 1C).

We next performed two genome-wide CRISPR screens to identify genes modulating LB-100 toxicity. Using HT-29 cells as a model, we carried out a CRISPRa screen to identify genes whose overexpression increases LB-100 toxicity. This screen identified 53 genes whose overexpression was selectively toxic in the presence of LB-100 (Figure S2A and Supplementary table S2). Among these genes we find proto-oncogenes often upregulated in human cancers (i.e., BCL9L, CCND1, CCNE2, ETS1, MAP3K1, MYC, MYB), suggesting that increased oncogenic signaling sensitizes cancer cells to LB-100 toxicity. In a complementary CRISPR-KO screen in SW-480 cells, we interrogated which gene knockouts could attenuate LB-100 toxicity. Figure S2B shows that gRNAs targeting genes from the WNT/β-catenin (CTNNB1, BCL9L, and LEF1) or MAPK (MAPK14/p38α, MAPK1/ERK2) signaling pathways were significantly enriched in the samples treated with LB-100 (Figure S2B and Supplementary table S3). These data indicate that suppression of WNT/β-catenin and MAPK signaling can alleviate the toxicity of LB-100. We combined the hits from both screens and built a String network ^18^ to unbiasedly infer cellular processes modulating LB-100 toxicity. Indicating only high-confidence interactions, the topology of the network supports the notion that oncogenic signaling modulates LB-100 toxicity (Figure 1D). Gene Ontology (GO) analyses found that terms associated with β-catenin pathway activity were among the top 3 enriched Biological Processes and Molecular Functions, while MAPK activity was the top enriched GO Molecular Function (Figure 1E). Together, these data indicate that activation of oncogenic signaling pathways lies at the heart of LB-100 toxicity in CRC cells. Positive modulation of the WNT/β-catenin and MAPK pathways increased LB-100 toxicity while losing components of these oncogenic pathways alleviated such toxicity.

### LB-100 is synthetic lethal with WEE1 inhibition

We then focused on the notion that increased oncogenic signaling intensifies the dependence on stress response pathways to support cancer cell viability^8^ (Figure 1 A, B). This raises the possibility that targeting of stress response pathways might be synergistic with further activation of oncogenic signaling in killing cancer cells. To investigate this, we designed a custom, stress-focused drug library comprised of 164 selected compounds targeting stress response pathways often associated with malignant phenotypes (i.e., DNA damage stress, oxidative stress, mitotic stress, proteotoxic stress, metabolic stress) (Supplementary table S4). The compounds were selected based on their ability to either induce these stresses or inhibit responses to them. Because senescence can be viewed as a survival response of cells under stress, we also included in the library senolytic drugs; drugs that selectively kill senescent cells^19^. Using both HT-29 and SW-480 cells as models, we tested each of these compounds at 15 concentrations, both in the presence and absence of a sub-lethal dose of LB-100 (2.5 µM) (Figure 2A). The differences in the normalized area under the curve (AUC) (with versus without LB-100) for each compound are represented in Figure 2B and Supplementary tables S5 and 6. We found that LB-100 increased the toxicity of inhibitors targeting CHK1 and WEE1 in both cell lines (i.e., CHK1 inhibitor GDC-0575 and WEE1 inhibitor adavosertib). Other inhibitors of these kinases were identified in one of two cell lines: CHK1i CCT-245737 and SCH-900776 in HT-29; and CHK1i rabusertib and prexasertib in SW-480. The dual CHK1/WEE1 inhibitor PD0166285 was also identified in SW-480 cells (Figure 2B). Using GDC-0575 as a CHK1 and adavosertib as a WEE1 inhibitor, we validated that adding LB-100 increases the toxicity of both drugs in these CRC cells (Figure 2C). Thus, the stress-focused drug screens identify CHK1 or WEE1 inhibition as a vulnerability of CRC cells treated with LB-100.

**Figure 2:**
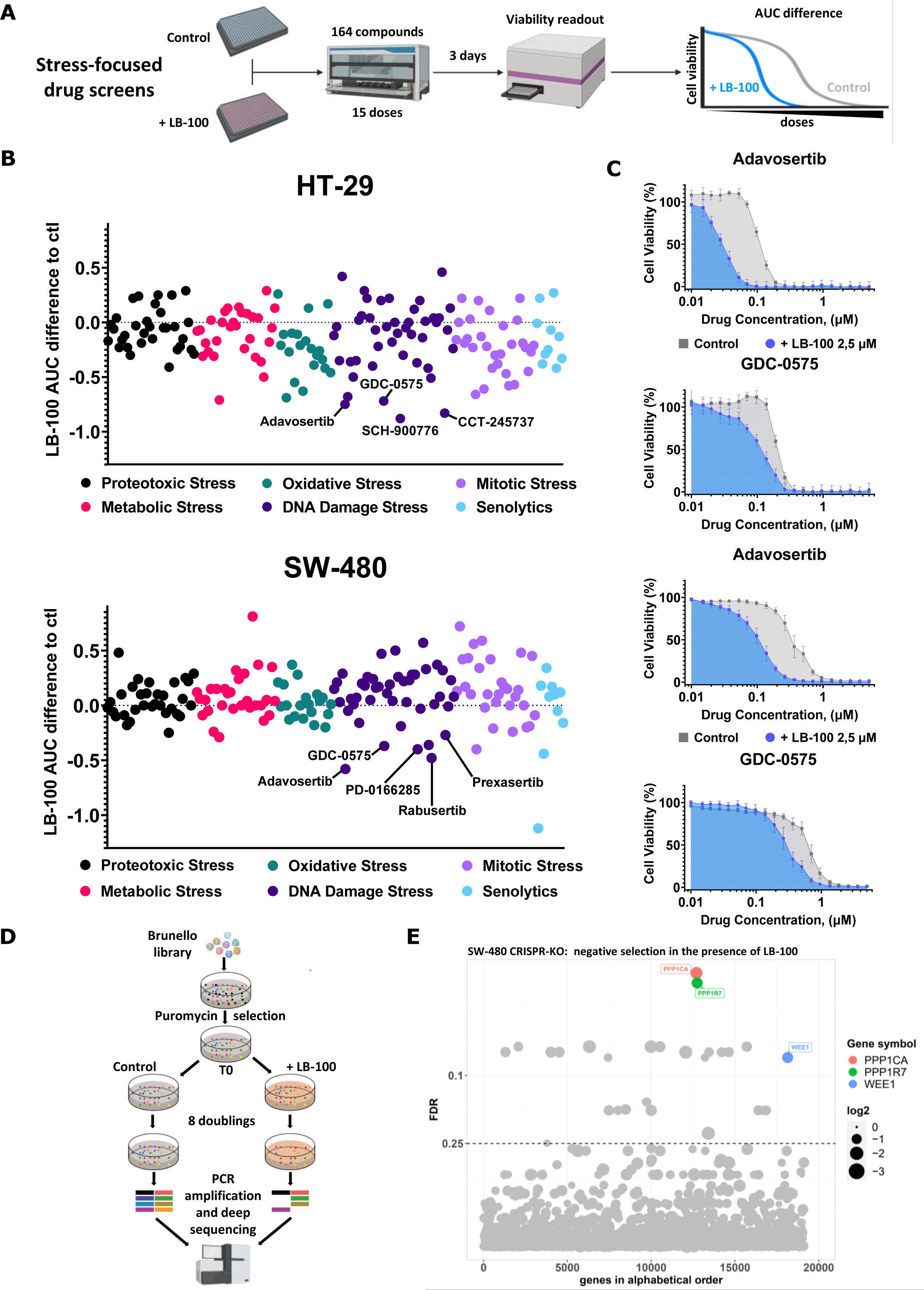
Stress-focused drug screen and genome-wide CRISPR screen converge to identify synthetic lethality between LB-100 and WEE1 inhibition. (A) Schematic outline of the stress-focused drug screen. **(B)** Area under the curve (AUC) difference for each compound in the presence of LB-100 (2.5 µM) relative to untreated controls in HT-29 and SW-480 cells. In both cases, WEE1 and CHK1 inhibitors are annotated. **(C)** Dose-response curves comparing the normalized AUC for Adavosertib or GDC-0575 in the presence or absence of LB-100 (2.5 µM) in HT-29 and SW-480 cells. Cell viability was estimated by resazurin fluorescence after 3 days in the presence of the drugs. **(D)** Schematic outline of the CRISPR-KO screen. **(E)** The bubble plot shows gRNAs significantly depleted in the LB-100 (2.5 µM) treated arm comparing to the untreated controls. 4 different gRNAs per gene were tested in 3 replicates. Cells on both conditions were grown for at least 8 population doublings before DNA harvesting and sequencing. Hits were called based on 0.25 false discovery rate (FDR) and at least 1 log2 fold-change difference between treated and untreated samples. Only the hits mentioned on the main text are named and colored, the full list of hits is presented on the Supplementary table S7 .

As an unbiased investigation of potential vulnerabilities of cells treated with LB-100, we performed another genome-wide CRISPR-KO screen in SW-480 cells looking for genes whose knockout is selectively toxic in the presence of the drug (Figure 2D). We found that gRNAs targeting 17 genes were significantly depleted in the LB-100 treated samples compared to the untreated controls (Figure 2E and Supplementary table S7). Among these genes are the catalytic (PPP1CA) and one regulatory subunit (PPP1R7) of Protein Phosphatase 1 (PP1), indicating an increased dependence on the activity of this other major Ser/Thr protein phosphatase^20^ upon PP2A inhibition. Interestingly, and consistent with the compound screens, gRNAs targeting WEE1 were also significantly depleted from LB-100 treated samples compared to the untreated controls (Figure 2E).

Data from the stress-focused drug screens and the genome-wide CRISPR screen converge to identify synthetic lethality between LB-100 and WEE1 inhibition in two CRC models. CHK1 inhibitors also showed increased toxicity in combination with LB-100 in our drug screens. However, the clinical development of most of these inhibitors has been discontinued^21^ discouraging further investigation. We therefore focused subsequent experiments on validating the combination of LB-100 and WEE1 inhibition in a panel of CRC models and a mechanistic understanding of the combined toxicity.

### LB-100 is synergistic with adavosertib in multiple cancer models

We used adavosertib as a WEE1 inhibitor to test the effect of the combination with LB-100 in the panel of seven CRC cell lines. Adavosertib dose-response assays indicated IC50s ranging from around 0.18 to 1 µM across the panel (Figure S3A). We then sought to address the toxicity of the combination in long-term viability assays (see methods). To that end, we first tested LB-100 and adavosertib as single drugs in this experimental setup and found variable toxicity of both drugs across the panel (Figure S3B), as anticipated by the drug-response assays. Informed by the toxicity of the single drugs, we addressed how sublethal concentrations of each drug would increase toxicity in combination. The results indicate strong toxicity of the combination using concentrations at which the single drugs show, at best, a modest effect (Figure 3A). It is noteworthy that DLD1, HCT-116, and SW-480 were largely tolerant to up to 500 nM of adavosertib, but such tolerance was abolished in the combination with LB-100 (Figure 3A). Toxicity of the combination was further confirmed across the CRC panel by IncuCyte-based cell proliferation assays. Doses of LB-100 or adavosertib that individually have a mild or transient impact ensued a more sustained restraint of cell proliferation in combination (Figure S3C). Next, we expanded the range of doses tested, to address if the two drugs act synergistically in these CRC cells. Synergy matrices combining 5 doses of each drug showed toxicities larger than expected based on the effect of the single drugs, as indicated by the respective synergy scores in five out of seven cell lines (Figure 3B), while DiFi and RKO cells showed synergy scores slightly below the proposed threshold of 10 for synergy^22^. These results confirm the combined toxicity of LB-100 and adavosertib in a diverse set of CRC cell lines, indicating no critical dependence on a specific set of oncogenic driver mutations in colorectal cancer.

**Figure 3:**
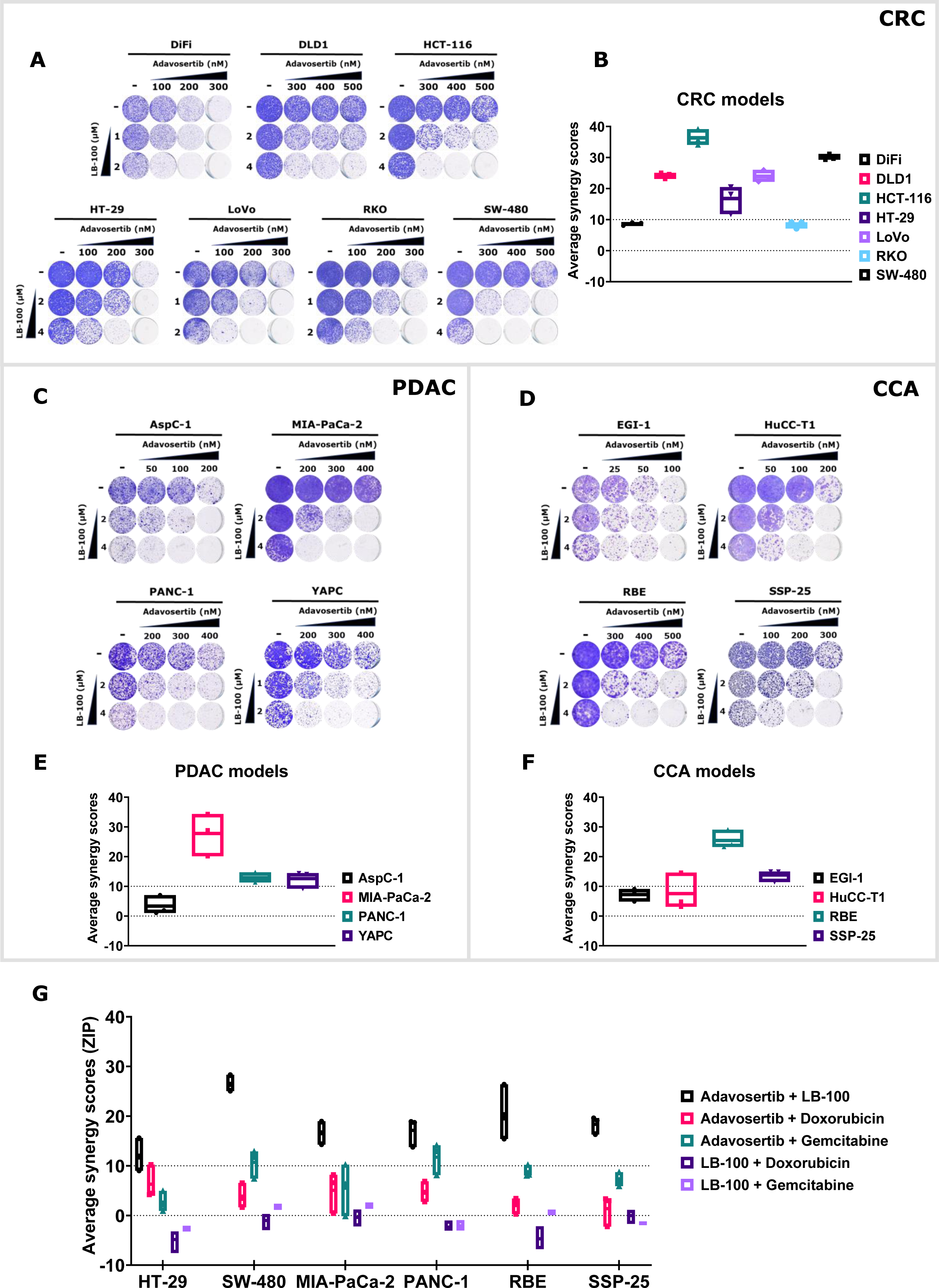
The combination of LB-100 and adavosertib is synergistic in cancer cells from different tissues and diverse genetic background. (A) Long-term viability assays show 7 CRC models treated with LB-100, Adavosertib, or the combination at the indicated concentrations. Cultures were refreshed every 2-3 days, and the cells were grown for 10-14 days before fixing, staining, and imaging. **(B)** Synergy scores for the combination of LB-100 and Adavosertib across 7 CRC models. Cells were treated with 5 concentrations of LB-100 (1, 2, 3, 4, and 5 µM) or Adavosertib (100, 200, 300, 400, and 500 nM) and all respective permutations for 4 days. The percentage of cell viability for each condition was estimated by resazurin fluorescence and normalized to DMSO controls. Synergyfinder.org web tool was used to calculate the ZIP synergy scores. 3 independent experiments are represented. **(C)** and **(D)** Long-term viability assays for 4 PDAC and 4 CCA models, respectively, treated with LB-100, Adavosertib, or the combination at the indicated concentrations. Cultures were refreshed every 2-3 days, and the cells were grown for 10-14 days before fixing, staining, and imaging. **(E)** and **(F)** Synergy scores for the combination of LB-100 and Adavosertib across 4 PDAC and 4 CCA models, respectively. Cells were treated with 5 concentrations of LB-100 (1, 2, 3, 4, and 5 µM) or Adavosertib (100, 200, 300, 400, and 500 nM) and all respective permutations for 4 days. The percentage of cell viability for each condition was estimated by resazurin fluorescence and normalized by DMSO controls. Synergyfinder.org web tool was used to calculate the ZIP synergy scores. 3 independent experiments are represented. **(G)** Synergy scores for the indicated combinations of LB-100, Adavosertib, Doxorubicin, and Gemcitabine across 6 cancer cell models. Cells were treated for 4 days with 10 concentrations of each drug: LB-100 (0.5, 1, 2, 3, 4, 5, 6, 7, 8, and 9 µM); Adavosertib (50, 100, 200, 300, 400, 500, 600, 700. 800, 900 nM); Doxorubicin (5, 10, 20, 30, 40, 50, 60, 70, 80, and 90 nM); Gemcitabine (0.63, 1.25, 2.5, 5, 10, 20, 30, 40, 50, and 60 nM); and all respective permutations for the combinations tested. The percentage of cell viability for each condition was estimated by resazurin fluorescence and normalized by DMSO controls. Synergyfinder.org web tool was used to calculate the ZIP synergy scores. 3 independent experiments are represented.

The efficacy of the combination in the CRC models encouraged us to test it in other tumor types lacking effective treatment options. Pancreatic ductal adenocarcinomas (PDACs) are refractory to conventional therapies and the 5-year survival rates remain one of the lowest among all cancers^23^. Similarly, despite a much lower overall incidence, cholangiocarcinomas (CCAs) share with PDACs the frequent lack of response to conventional therapies and the dismal prognosis^24^. We put together a small panel of four PDAC cell lines, and a similar one of CCA cells (Table S1) to assess the potential of the combination of LB-100 and adavosertib in these cancer types. LB-100 dose-response assays revealed IC50s varying from 3.2 to >10 µM in the PDAC cells (Figure S4A), and from 4.7 to >12 µM in the CCA models (Figure S4D). For adavosertib, the IC50s ranged from 0.3 to 1.7 µM in the PDAC models (Figure S4A), and from 0.1 to 0.37 µM in the CCA models (Figure S4D). To test the effects of the combination in the PDAC and CCA cancer models systematically, we employed the same experimental workflow used for the CRC panel. We first addressed the long-term toxicity of LB-100 and adavosertib in the PDAC and CAA models (Figures S4B and E). Then, we combined sublethal doses of each drug and found strong or complete suppression of cell viability in the models from both cancer types (Figures 3C and D). Cell proliferation assays corroborate that doses of the individual drugs that are ineffective in suppression of cell proliferation in each of the PDAC and CCA models become quite effective when used in combination (Figures S4C and F). Furthermore, matrices combining LB-100 and adavosertib indicated synergy in 3 out of the 4 PDAC models (Figure 3E). For the CCA panel, we found synergy scores above the threshold for RBE and SSP-25, but not for EGI and HuCC-T1 cells (Figure 3F).

Pre-clinical data support the efficacy of adavosertib in combination with chemotherapy, which has subsequently been investigated in multiple clinical trials^25^. Likewise, LB-100 has been proposed as a sensitizer for chemotherapy^14^ and is currently under clinical investigation in such combination (NCT04560972). We asked how the combination of LB-100 and adavosertib compares to each of these drugs in combination with doxorubicin or gemcitabine. We performed synergy matrices using 10 concentrations of each drug in 2 cell models per tumor type. The results showed higher synergy scores for the combination LB-100 + adavosertib compared to combinations with the chemotherapeutic agents in each of the models (Figure 3G).

These data reveal considerable context independence of the synthetic lethality between LB-100 and adavosertib in cancer cell lines from different tissues and diverse genetic backgrounds. Moreover, they suggest that the combination proposed here might provide therapeutic benefits superior to regimens that are currently under clinical investigation. We, therefore, focused next on the mechanistic understanding of the toxicity and evaluating the viability of this combination *in vivo*.

### The combination LB-100 and adavosertib leads to aberrant mitoses and cell death

WEE1 inhibition abrogates the G2 to M cell cycle checkpoint, allowing cells to enter mitosis without resolving DNA damage, leading to mitotic catastrophe and apoptosis^26^. Similarly, LB-100 has been shown to induce mitotic catastrophe in several tumor models, particularly in combination with DNA-damaging agents^15^. It is well-established that the unscheduled proliferation inherent to cancer cells comes at the expense of abnormal mitoses and associated mitotic stress^27^. This raises the question whether the observed synergy of these drugs results from fatal mitotic defects.

To address this, we followed HT-29 cells expressing H2B-GFP by live-cell imaging from nuclear envelope breakdown (marking mitotic entry) until anaphase onset. Numerous positive feedback loops ensure complete commitment to mitosis upon mitotic entry^28^, and such an “all-in” process takes place in a fraction of the cell cycle time. Accordingly, we found that DMSO-treated cells spent an average of 50 minutes in mitosis. LB-100-treated cells, however, spent on average over 600 minutes in mitosis, with prominent variability among individual cells. Adavosertib also increased the average time spent in mitosis, although to a lesser extent (∼100 minutes). The combination resulted in an average time in mitosis of over 1050 minutes with virtually all cells showing markedly long mitoses (Figure 4A). Detailed analyses of the time-lapse images revealed that defective chromosome alignment is highly prevalent in the LB-100-treated cells in the first hours after the addition of the drug, which was alleviated in cells entering mitosis later (Figure 4C, 2^nd^ panel). Adavosertib induced fewer but similar misalignments in cells entering mitosis later and sharply reduced the average time for mitotic entry to less than 6h, compared to about 10h in the control and LB-100-treated cells (Figure 4C, 3^rd^ panel). Cells treated with the combination showed both the accelerated mitotic entry observed for adavosertib and the high prevalence of misalignments shortly after drug exposure observed for LB-100. Strikingly, most of the combination-treated cells entering mitosis later showed catastrophic mitoses with only partial chromatin condensation and failed attempts to divide until the end of the experiment (Figure 4C 4^th^ panel). This phenotype was not observed in the cells treated by the single drugs. These results reveal that combination therapy in HT-29 CRC cells results in a defective mitotic-like state that persists for at least 24h following drug exposure. Although the time-lapse experiments did not indicate the fate of these cells, measurements of caspase 3/7 activity for 96h show apoptosis induction in the cells treated with the combination (Figure 4D).

**Figure 4:**
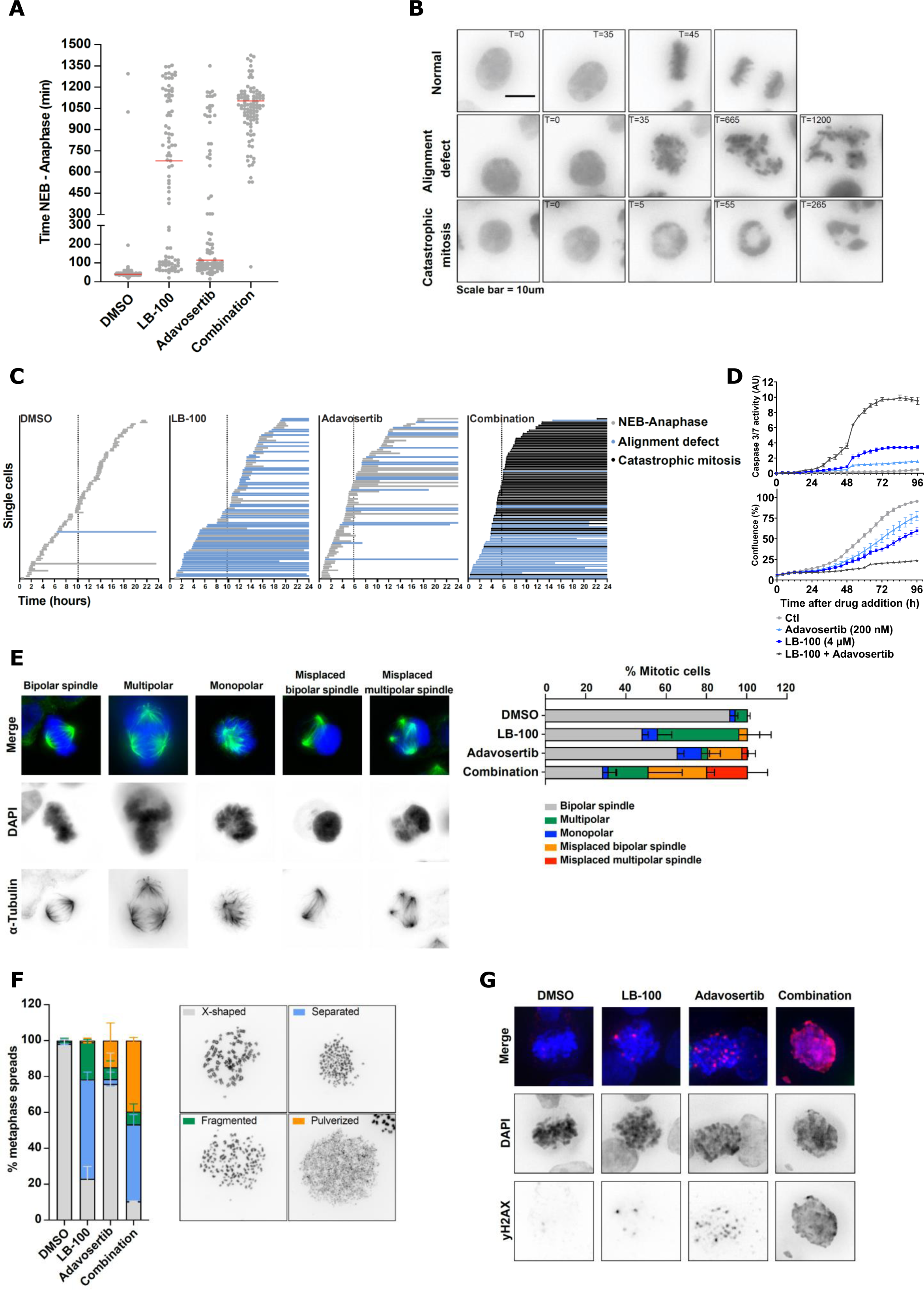
LB-100 and adavosertib combination leads to aberrant mitoses and cell death. (A) Time from nuclear envelope breakdown (NEB) to anaphase for HT-29 cells untreated (DMSO) or treated with LB-100, Adavosertib, or the combination. Each dot represents an individual cell followed by live-cell imaging. Red bars represent the average time spent from NEB to anaphase. 2 independent experiments are compiled (n = 100 cells per condition). **(B)** Representative live-cell microscopy images of HT-29 cells. The examples highlight the two major mitotic phenotypes observed. The scale bar represents 10 µm. **(C)** Representation of the time for mitotic entry and exit of HT-29 cells imaged every 5 minutes for 24h, starting immediately after the addition of DMSO, LB-100, Adavosertib, or the combination. Each bar represents an individual cell. The colors of the bars indicate normal or aberrant mitoses. The beginning of the bars mark NEB and the end represents either anaphase or end of the experiment. Dashed vertical lines represent average times for mitotic entry after the addition of the drugs. **(D)** IncuCyte-based assay for caspase 3/7 activity. Cells were treated with DMSO, LB-100, Adavosertib, or the combination in the presence of a caspase 3/7 apoptosis assay reagent. Green fluorescence from the apoptosis assay reagent divided by the total confluence was used to estimate apoptosis for 96h. **(E)** Representative images (left) and quantification (right) of spindle defects in mitotic cells treated with DMSO, LB-100, Adavosertib, or the combination. Cells were treated for 8h before fixation. DNA was stained with DAPI (blue) and α-Tubulin was immunostained (green). Quantification is based in 2 independent experiments each analyzing 50 cells per condition **(F)** Chromosome spreads from HT-29 cells treated with DMSO, LB-100, Adavosertib, or the combination. On the left, quantification of chromosome integrity; on the right, representative images. Drugs were added for 16h then nocodazol was added for additional 3h to block cells in mitosis. Cells were harvested by mitotic shake-off for spreading. **(G)** Representative images show HT-29 cells treated with DMSO, LB-100, Adavosertib, or the combination for 24 h. After fixation, total DNA was stained with DAPI (blue) and γ-H2AX was immunostained (red). Throughout Figure 4, LB-100 was used at 4 µM and Adavosertib at 200 nM.

To better understand the observed mitotic defects, we next examined the microtubule organization of mitotic cells in response to the single drugs and the combination. The vast majority of the DMSO-treated cells showed the expected bipolar spindles attached to the metaphase plate. LB-100 treatment induced multipolar spindles in about 40% of the mitotic cells, while adavosertib increased the frequency of monopolar and misplaced bipolar spindles. Aberrant spindles were found in over 70% of the cells treated by the combination of these drugs (Figure 4E). The abnormalities observed in these mitoses also included multipolar and misplaced bipolar spindles. However, misplaced multipolar spindles were only frequent in combination-treated cells, further indicating mitotic defects emerging from the combination.

As an additional readout for mitotic defects, we addressed chromosome integrity across the different treatments. Chromosome spreads from DMSO-treated cells showed the expected paired chromatids attached at the primary constriction (Figure 4F). LB-100 treatment largely increased the frequency of separated and fragmented chromatids, which is consistent with the misalignment phenotype observed in Figure 4C. Adavosertib treatment compromised chromosome integrity in nearly 30% of the cells, with a fraction of them failing to produce discrete chromosomes. Such “pulverized” spreads were present in about 40% of the cells exposed to the combination, and separated chromatids, as observed for LB-100 single treatment, were also frequent (Figure 4F). These observations align with the misplaced spindles shown in figure 4E, given that those pulverized chromosomes could not establish a proper metaphase plate. Such an apparent chromosome fragmentation also indicates severe DNA damage in these mitotic cells. In line with that, imaging of mitotic HT-29 cells treated with the combination revealed a pan-nuclear γ-H2AX staining instead of punctate foci observed in LB-100- or Adavosertib-treated γ-H2AX-positive cells (Figure 4G). Altogether, these data indicate that the combined induction of catastrophic mitoses followed by cell death underlies the synergy between LB-100 and WEE1 inhibition in CRC cells.

### LB-100 and adavosertib cause concerted DNA replication stress, priming cancer cells to premature and deadly mitoses

It has been shown previously that WEE1 inhibition not only abrogates the G2/M checkpoint but can also drive S-phase cells under replication stress into premature mitoses^29^. Such a scenario would be in line with the aberrant mitoses with pan-nuclear DNA damage described above. We therefore asked whether and how replication stress contributes to the toxicity of the combination. The presence of single-strand DNA (ssDNA) resulting from uncoupled DNA helicase and polymerase activities is direct evidence of replication stress^30^. We used BrdU detection under native (non-denaturing) conditions as an indication of replication stress. We found that both LB-100 and adavosertib increased the percentage of HT-29 cells with multiple ssDNA foci 8h after drug exposure (Figure 5A). Even more prominent replication stress was present under the combined treatment, with around 50% of the cells showing multiple ssDNA foci (Figure 5A). Similar results were observed in SW-480 cells; however, adavosertib led only to a mild increase in ssDNA foci in this model (Figure 5B). We next performed DNA fiber assays on both CRC models to address the impact of these drugs on the dynamics of DNA replication. LB-100 had no significant effect on the average DNA replication fork speed in HT-29 and modestly increased it in SW-480 cells compared to untreated samples (Figures 5C and D). The percentage of replication origin firing in both cell models was similar to the respective controls. Adavosertib, however, led to a reduced average fork speed, as previously described^31, 32^, and a concomitant increase in the percentage of origin firing in both CRC models (Figures 5C and D). An inverse correlation between fork speed and origin firing is characteristic of DNA replication under perturbations^33^. This is accomplished by the usage of backup replication origins under stress, given that these origins are licensed in large excess relative to the amount necessary to complete an unperturbed S phase^34, 35^. It is therefore noteworthy that the concomitant addition of LB-100 further reduced the slower fork speed observed for adavosertib, but without the concurrent increase in origin firing (Figures 5C and D). Such a “decoupling” of the inverse correlation between fork speed and origin firing indicates complementary effects of these drugs to disrupt the dynamics of DNA replication forks in these CRC cells.

**Figure 5:**
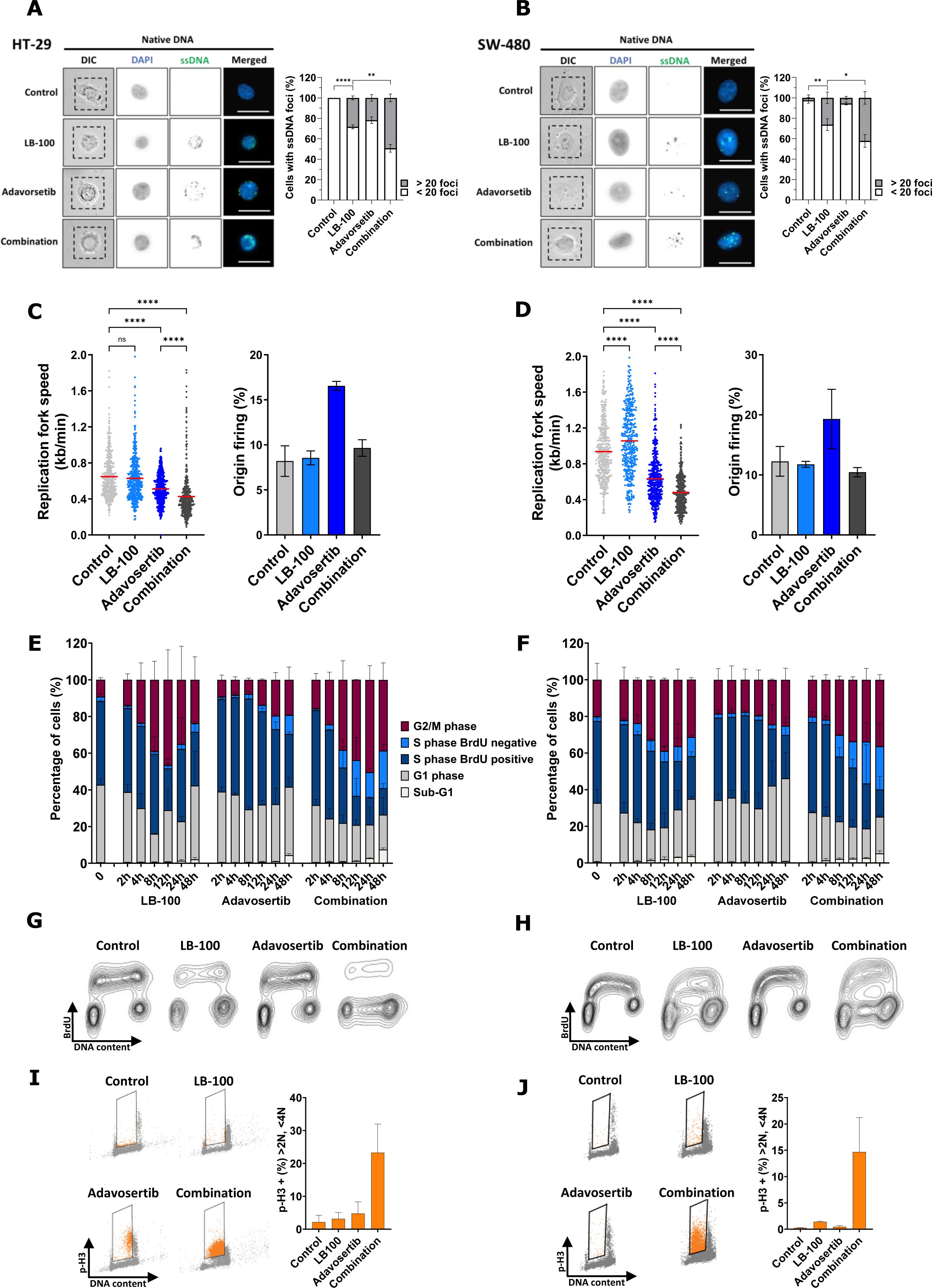
LB-100 and adavosertib promote concerted replication stress, priming for premature mitoses. Representative images and quantifications of single-strand DNA (ssDNA) foci in HT-29 **(A)** and SW-480 **(B)** cells. After incorporating BrdU for 48h, the cells were treated as indicated for 8h and then fixed. Total DNA was stained with DAPI (blue) and BrdU was immunostained (green) under non-denaturing conditions to indicate long fragments of ssDNA. Quantification is based on 2 independent experiments analyzing at least 100 cells per coverslip. Asterisks indicate significance level (* p-value <0.05, ** p-value <0.01, **** p-value <0.0001) by two-tailed unpaired t test. **(C)** and **(D)** DNA fiber assays show replication fork speed (left) and percentage of origin firing (right) of HT-29 and SW-480 cells, respectively, untreated (DMSO) or treated with LB-100, Adavosertib, or the combination for 8h. For fork speed, track lengths of at least 380 ongoing forks from each condition were measured with ImageJ in 2 independent experiments that are shown combined. Red lines indicate the mean and asterisks indicate significance level (**** p-value <0.0001) by nonparametric Kruskal–Wallis test. For origin firing, first label and second label origins are shown as percentage of all labelled tracks. At least 1000 labelled tracks per condition were analyzed. **(E)** and **(F)** Quantification of time-course cell cycle flow cytometry from HT-29 and SW-480 cells, respectively, treated with LB-100, Adavosertib, or the combination. Cells were fixed at the indicated time points after addition of the drugs. BrdU (10 µM) was added 1h before fixation. Total DNA was stained by Propidium Iodide (PI) and BrdU was immunostained. Cell cycle phases were gated using the FlowJo software. Error bars represent the standard deviation of 2 independent experiments. **(G)** and **(H)** Representative image of the 12h time point from the time-course flow cytometry above. **(I)** and **(J)** Flow cytometry assessment of p-H3 (Ser10) vs DNA content in HT-29 and SW-480 cells, respectively, treated with LB-100, Adavosertib, or the combination for 12h. Total DNA was stained by Propidium Iodide (PI) and p-H3 (Ser10) was immunostained. S-phase cells were gated using the FlowJo software. Error bars represent the standard deviation of 2 independent experiments. Throughout figure 5, LB-100 was used at 4 µM on both cell lines. Adavosertib was used at 200 nM in HT-29 and 400 nM in SW-480.

Next, we addressed how such a concerted action on replication forks impacts the cell cycle progression of these CRC cells. For that, we performed time-course flow cytometry analyses assessing total DNA content and BrdU incorporation as a proxy for active DNA synthesis. The results for both HT-29 (Figure 5E) and SW-480 models (Figure 5F) show a transient accumulation of cells in G2/M induced by LB-100, peaking around 12h after stimulation. More sustained G2/M accumulation was observed in the combination-treated samples, while adavosertib mildly increased the G2/M fraction (Figures 5E and F). Strikingly, in both CRC models, the samples treated with the combination showed an increased accumulation of cells with S-phase DNA content that are negative for BrdU, indicating a lack of active DNA synthesis (Figures 5E-H). These data confirm that the combination of LB-100 and adavosertib halts DNA replication, as indicated by the fiber assays. In fact, DNA replication is incompatible with a mitotic state, and stalled replication forks are actively cleaved in mitosis^36^. The lack of active replication and the pulverized chromosome spreads support a scenario in which the combination of LB-100 and adavosertib drives these CRC cells to mitosis before completing DNA replication. Corroborating this rationale, in both CRC models, cells with DNA content between G1 and G2/M (compatible with S phase) become positive for the mitosis marker phospho-histone H3 (p-H3) (Ser10) 12h after treatment with the combination (Figures 5I and J). Together, the data above indicate that the combination of LB-100 and adavosertib results in severe replication stress, priming these CRC cells to catastrophic mitoses prior to DNA replication completion.

### LB-100 and adavosertib combination restrains tumor growth *in vivo*

The data above show the efficacy of the LB-100 and adavosertib combination in multiple cancer models and how these compounds cooperate to disrupt cancer cell viability. Although the observed synergistic activity and context independence *in vitro* are promising, the balance between efficacy and toxicity must be tested *in vivo* to indicate a potential therapeutic window. We implanted tumor samples derived from three different CRC patients in the colon of immunodeficient mice to ask whether the combination LB-100 and adavosertib suppresses tumor growth *in situ*. It is noteworthy that these orthotopic patient-derived xenografts (PDXs) derive from metastatic colorectal tumors with diverse mutation backgrounds that progressed under previous treatment regimens (see methods). After mice randomization, we treated these PDXs with the single drugs and the combination for four weeks and measured endpoint tumor sizes at sacrifice. The results showed antitumor activity of LB-100 in two of the three PDXs, while adavosertib restrained tumor growth in all three PDXs. Yet, in combination, these drugs strongly suppressed tumor growth, showing antitumor activity significantly superior to the single drugs in 2 of the 3 PDXs (Figure 6A).

**Figure 6:**
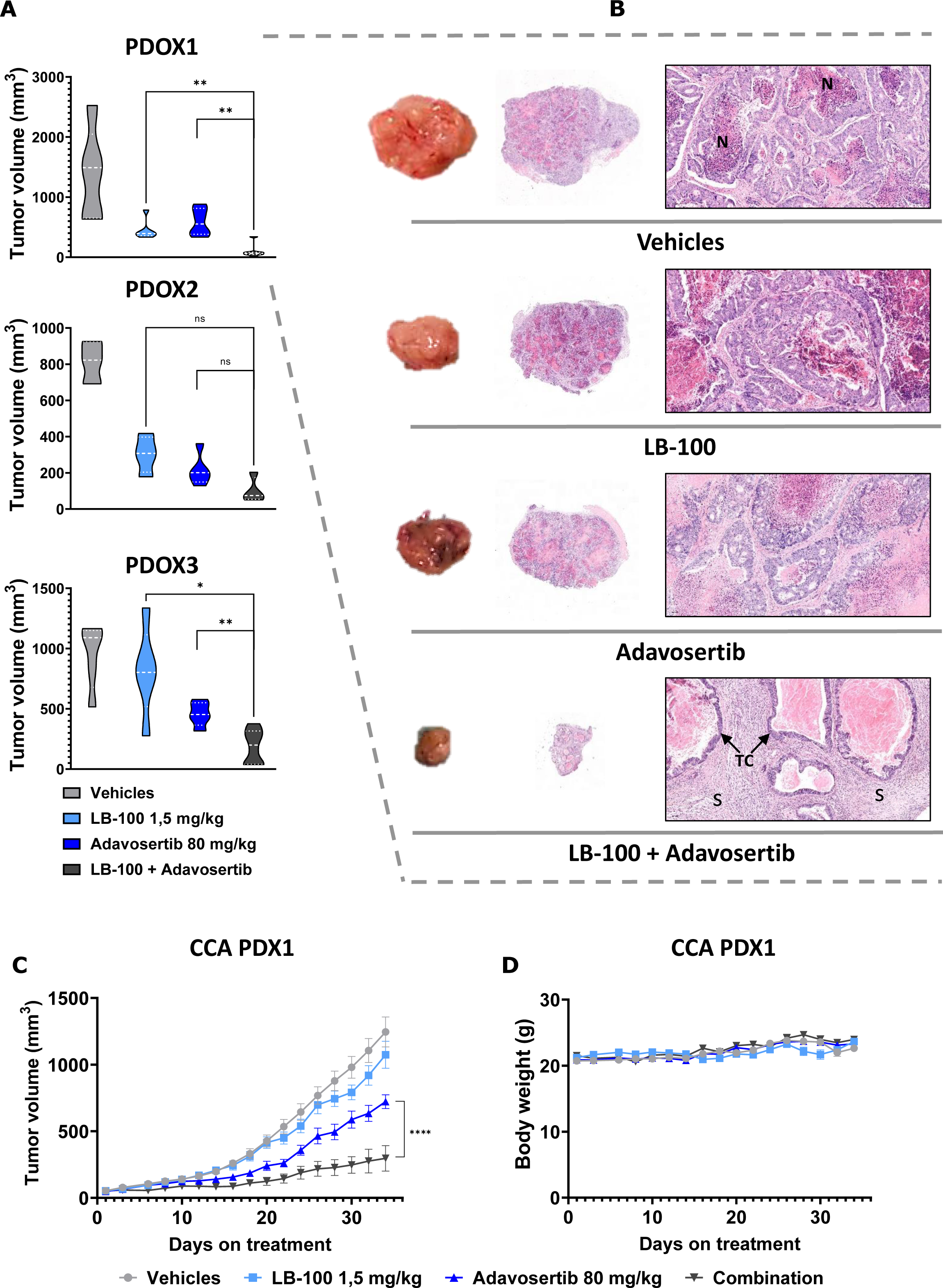
LB-100 and adavosertib combination restrains tumor growth *in vivo*. (A) Endpoint tumor volumes of 3 independent orthotopic CRC PDXs treated with LB-100, adavosertib, or the combination for 4 weeks. After transplantation and engraftment, the mice were randomized and treated as indicated. LB-100 was given on days 1, 3, and 5, while Adavosertib was administered on days 1-5 in 7-day cycles . Asterisks indicate significance level (* p-value <0.05, ** p-value <0.01) by two-tailed Mann-Whitney test. **(B)** Representative tumors and Hematoxylin & Eosin stainings at endpoint from PDOX1 treated as indicated. Original magnification middle images: 15x, scale bar 1000 µm; right images: 200x, scale bar 100 µm. N indicates necrotic areas, S indicates stroma, and the arrows point to the tumor-cell component. **(C)** Tumor growth curves of cholangiocarcinoma PDX1 treated with LB-100, adavosertib, or the combination. After transplantation and engraftment, the mice were randomized and treated as indicated. LB-100 was given on day 1, adavosertib was given on days 1-3, in 4-days cycles. Tumors were measured 3 times per week. The graphs show mean and SEM. Asterisks indicate significance level (**** p-value <0.0001) by two-way ANOVA with Tukey’s multiple comparisons. **(D)** Body weight variation of the CCA PDX1 across the experiment. The graphs show mean and SEM.

In addition to the reduction of tumor size, histopathological features of tumor regression indicate treatment efficacy and predict patients’ prognoses^37^. We therefore addressed the endpoint histology of these patient-derived tumors under the four treatment regimens. As highlighted in Figure 6B, vehicle-treated PDOX1 tumors showed moderately differentiated adenocarcinoma, consisting of closely packed glands with “dirty” luminal necrosis. No obvious histopathological regression was observed in the tumors treated with LB-100 or adavosertib. However, clear histopathological regression was found in tumors treated with the combination, showing reduced tumor-cell component, and increased fibrosis and inflammatory infiltrate (Figure 6B). The moderately differentiated adenocarcinoma PDOX2 showed similar histological regression for the combination and single drugs (Figure S5A left). The histology of the poorly differentiated adenocarcinoma PDOX3 tumors was in line with the observed for PDOX1, with similar histopathological responses for the combination and no obvious effect of the single drugs (Figure S5A right). The body weight curves indicate that the single drugs and the combination were well tolerated in these mouse models (Figure S5B). However, 1/15 mice treated with LB-100 and 2/22 treated with the combination died during the experiment. These casualties cannot be unequivocally attributed to treatment toxicity (P values >0.45). Moreover, we dosed LB-100 at 1.5 mg/kg; this drug has been extensively tested in multiple mice models in doses ranging from 1.5 to 2.5 mg/kg without noticeable toxicity, even in combination with chemotherapy^14^.

Expanding our analyses to a different tumor type, we transplanted patient-derived cholangiocarcinoma tumor fragments into the flank of immunocompromised mice. After engraftment and randomization, we treated these mice with LB-100, adavosertib, or the combination until the vehicle-treated control group reached the endpoint. In this PDX, LB-100 alone showed little antitumor effect, while adavosertib delayed tumor growth more obviously. Yet, the combination promoted an antitumor effect clearly superior to any of the single drugs and strongly restrained tumor growth in this CCA PDX (Figure 6C). It is noteworthy that no casualties were observed in any treatment groups throughout the experiments and the body weight curves indicate that the combination was tolerated by the mice (Figure 6D). These results confirm the antitumor activity of the combination LB-100 + adavosertib in patient-derived models using doses not associated with systemic toxicity.

### Acquired resistance to the combination of LB-100 and adavosertib is tumor-suppressive

Even highly synergistic drug combinations can ultimately result in resistance in patients with advanced disease^38^. Since deliberate further activation of oncogenic signaling fundamentally differs from inhibition of these signals, we studied how cancer cells may acquire resistance to the combination of LB-100 and adavosertib. We selected polyclonal pools of HT-29 and SW-480 resistant cells (HT-29-R and SW-480-R) by culturing parental cells in the presence of the drug combination for over four months. Long-term viability assays illustrate the reduced toxicity of the combination in the resistant cells compared to the respective parental cells (Figure 7A). IncuCyte-based proliferation assays showed that both combination-resistant models proliferate slower than their respective parental cells (Figure S6A). Moreover, despite growing in the presence of the drugs for several months, there is no apparent "addiction" to the combination. Instead, for HT-29 combination-resistant cells, the combination still hinders cell proliferation (Figures S6A left).

**Figure 7:**
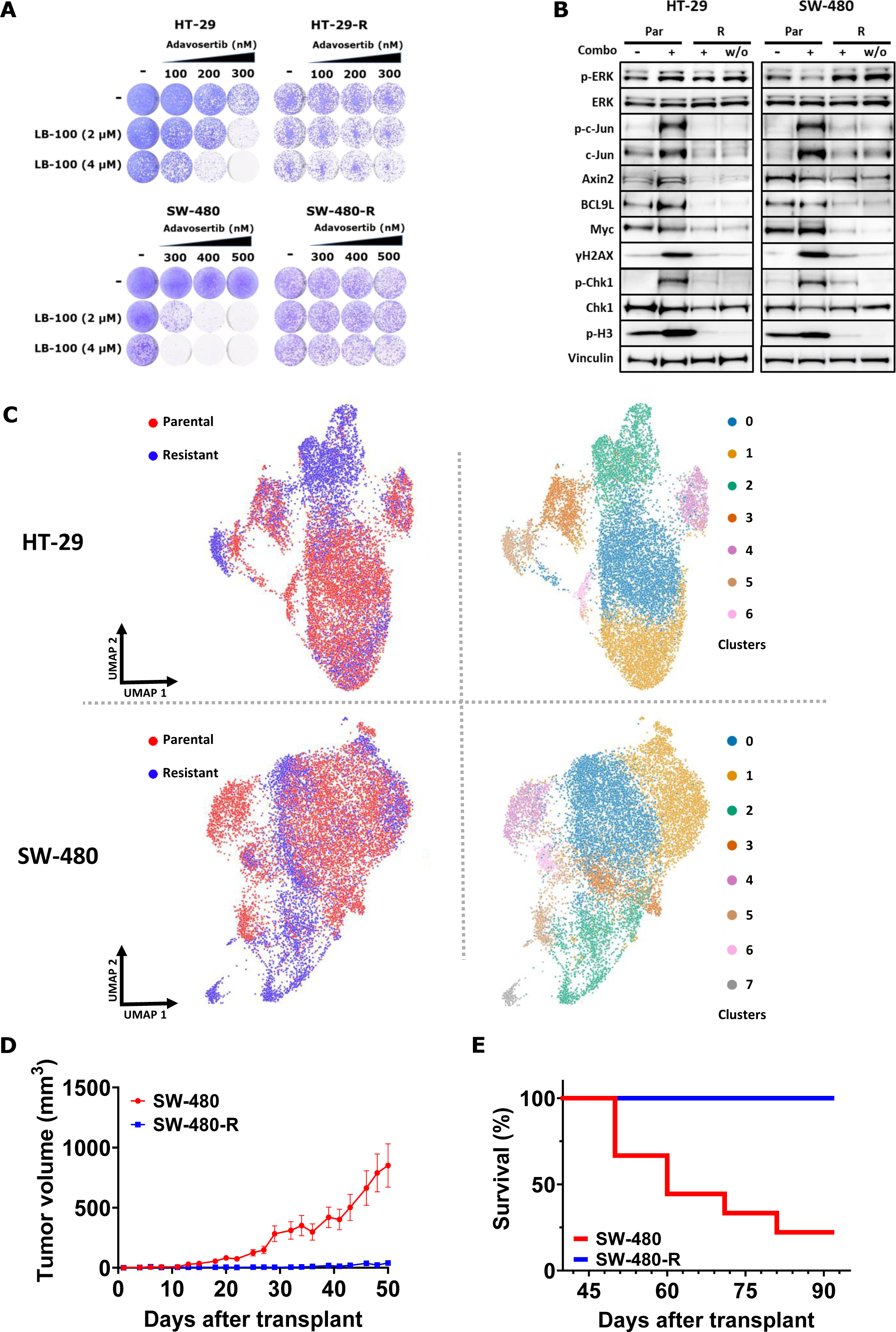
Acquired resistance to the combination of LB-100 and adavosertib is tumor-suppressive. (A) Long-term viability assays show HT-29 and SW-480 parental and resistant cells treated with LB-100, Adavosertib, or the combination at the indicated concentrations. Cultures were refreshed every 2-3 days, and the cells were grown for 10-14 days before fixing, staining, and imaging. **(B)** Western blots show selected oncogenic signaling and stress response pathways in HT-29 and SW-480 parental and resistant cells. Parental cells were exposed to the combination for 24h, while for resistant cells, that grow in the presence of the combination, the drugs were washed out (w/o) 24h before harvesting. LB-100 was used at 4 µM and adavosertib was used at 400 nM. Vinculin was used as loading control. **(C)** UMAP representations of HT-29 (top) and SW-480 (bottom) parental and resistant cells colored by sample of origin (left) or by clusters (right). Parental cells were harvested untreated and resistant cells were harvested 24h after the washout of the drugs. **(D)** Tumor growth curves of SW-480 parental and resistant cells in the absence of drugs. Per cell line, 10 mice were injected subcutaneously with 3 million cells and we measured tumor size 3 times per week. The graph shows mean and SEM of the measurements until the first mouse reached 1500 mm^3^ (ethical sacrifice). **(E)** Kaplan-Meier survival curves of the experiment above along 3 months considering the 1500 mm^3^ ethical sacrifice.

We reasoned that acquired resistance to hyperactivation of oncogenic pathways might lead to the down-modulation of oncogenic signaling to evade the stress imposed by the increased signaling. Indeed, c-JUN is no longer hyperactivated in the resistant cells in the presence of drugs, suggesting downmodulation of this MAPK signaling arm. Furthermore, the levels of the β-catenin targets AXIN2 and MYC and the modulator of β-catenin transcriptional activity BCL9L were lower in the resistant cells, even in the absence of the drugs (Figure 7B). In contrast, p-ERK levels remained higher in the combination-resistant cells compared to parental controls. For both CRC models, p-CHK1, γ-H2AX, and p-H3 (Ser10) levels are also no longer increased in the presence of the drug (Figure 7B). These data indicate down-modulated β-catenin and JUN signaling coincident with alleviated replication and mitotic stresses after acquired resistance to LB-100 + adavosertib in these CRC cells.

Our data show the presence of numerous mitotic defects induced by the combination of LB-100 and adavosertib in these CRC cells (Figure 4). HT-29 and SW-480 karyotypes have been described as near triploid^39^, illustrating the aneuploidy characteristic of cancer cells. We asked whether acquired resistance to that combination impacted the aneuploidy in these CRC models. To address this, we prepared chromosome spreads from the parental and combination-resistant cells and counted the number of chromosomes. The results showed that HT-29-R cells have a median chromosome number of 62, compared to about 66 on HT-29 parental cells (Figure S6B left). SW-480 parental cells showed marked heterogeneity in the number of chromosomes across the population, with a median number of 80. Such heterogeneity was sharply reduced in the SW-480-R cells that showed a median of 54 chromosomes (Figure S6B right). These data evidence reductions of aneuploidy during the acquired resistance to the combination of LB-100 and adavosertib in these CRC cells.

To gain a deeper understanding of the nature of the resistance and the heterogeneity of these polyclonal populations, we performed single-cell RNA-seq on HT-29-R and SW-480-R and their respective parental cells. We analyzed over 4000 cells per model, which yielded 6 UMAP clusters for HT-29 and 7 for SW-480 cells after processing and integration, performed independently per cell line (see methods). Interestingly, for HT-29, we can observe that cluster 2 is virtually absent in parental cells, while they compose most of the clusters 0 and 1 (Figure 7D top). For SW-480 cells, while clusters 2 and 7 are populated mostly by resistant cells, the major clusters 0 and 1 show a mix of parental and resistant cells (Figure 7D bottom). By analyzing the marker genes making the identity of each cluster, we observe that, for HT-29 cells, clusters 0 and 1 are defined by several genes associated with proliferation (e.g., MCM3/4/6, CDC6, PLK1, AURKA, CCNB1/2, CCNA2, MKI67, among others). The resistance-associated cluster 2 shows a rather low expression of most of these genes and displays marker genes associated with inflammation (Figure S6C left). In SW-480 cells, although the specific genes vary, similar inflammation-related markers were observed for cluster 2, which is also composed of combination-resistant cells. In this model, only a few markers discriminate cluster 0 from the other cells. Cluster 1, however, showed expression enriched for genes associated with proliferation, resembling clusters 0 and 1 from HT-29 (Figure S6C right). Because our data indicate that β-catenin and MAPK signaling outputs modulate sensitivity to LB-100 (Figure 1D and S2), we also computed the activity of these pathways at the single-cell level in parental and combination-resistant cells (see methods). No clear difference in the MAPK pathway activity scores was observed between parental and resistant HT-29 cells (Figure S7A top left), while SW-480-R scored slightly higher than SW-480 parental cells (Figure S7B top left). On the other hand, β-catenin pathway scores were lower in the resistant cells from both models (Figure S7A and B top right). Furthermore, both MAPK and β- catenin pathways modulate the transcriptional activity of MYC^40, 41^. We can observe on the UMAPs that lower MYC targets scores mostly correspond to combination-resistant cells (Figure S7A and B bottom left), indicating a decreased MYC activity. Likewise, we found lower E2F targets pathway activity scores on HT-29 and SW-480 cells that acquired resistance to LB-100 + adavosertib (Figure S7A and B bottom right). This is noteworthy because elevated expression of E2F targets is frequent in tumors, and higher levels have been associated with poor prognosis in different cancers^42^. Overall, the single-cell RNAseq data support the notion that acquired resistance to the drug combination results in reduced mitogenic signaling and attenuation of the proliferative transcriptional program characteristic of cancer cells.

The data above showing downmodulation of oncogenic signaling, reduced aneuploidy, and transcriptional shifts associated with less proliferative phenotypes suggest an intriguing outcome for the acquired resistance to the combination of LB-100 and adavosertib: suppression of the malignant phenotype. Anchorage-independent proliferation is a common trait of transformed cells and can be used as a proxy for malignancy. Therefore, we asked how acquired resistance to this combination would impact anchorage-independent proliferation in these CRC models in the absence of the drugs. Parental SW-480 cells showed similar endpoint viability both in attached or anchorage-independent conditions. Strikingly, the anchorage-independent proliferation of SW-480-R cells was around 10-fold lower than measured for the parental cells. (Figure S8A right). Similar results were observed in HT-29 cells. However, these cells showed a lower anchorage-independent proliferation, compressing the comparative difference between parental and resistant cells (Figure S8A left). Finally, we directly addressed the notion of reduced tumorigenicity associated with the acquired resistance. We transplanted SW-480 parental (which showed better anchorage-independent growth) and SW-480-R cells into immunocompromised mice and monitored tumor growth. The results showed clear engraftment within the first 25 days and steady tumor growth in mice transplanted with SW-480 parental cells. Conversely, SW-480-R cells either failed to develop or developed small tumors over 50 days after transplantation (Figure 7D). The mice were sacrificed at the ethical endpoint of 1500 mm^3^. Kaplan-Meier survival curves show that none of the ten tumors from mice injected with SW-480-R cells reached that endpoint during the 3 months of the experiment. In contrast, 7 out of 9 tumors from parental SW-480 cells reached the 1500 mm^3^ endpoint during the experiment (Figure 7E). Spider plots further illustrate the poor engraftment of the combination-resistant cells compared to the parentals, despite evident heterogeneity on the growth kinetics of the individual tumors (Figure S8B).

Together, these data indicate that chronic exposure to the LB-100 and adavosertib combination may lead to acquired resistance that is fundamentally distinct from what is seen with drugs that inhibit oncogenic signaling: suppression of the malignant phenotype.

## DISCUSSION

We explored here an unconventional rationale for cancer therapy based on further activation of oncogenic signaling rather than trying to inhibit it. This approach is inspired by mounting experimental data indicating that the homeostasis of cancer cells relies on optimal levels of oncogenic signaling, not necessarily the highest level ^4–6, 9^. To support the malignant state, stress response pathways act as buffers to the detrimental effects of oncogenic signaling^8, 9^. In the present study, we used the PP2A inhibitor LB-100 to hyperactivate oncogenic signaling, leading to multiple stresses in CRC cells. Using both compound and genetic screens, we found that concomitant inhibition of the mitotic gatekeeper kinase WEE1 is lethal in multiple cancer models, and this combination restrained the growth of patient-derived tumors in mice.

That deliberate overactivation of oncogenic signaling is toxic to cancer cells is not without precedent. For instance, pharmacological upregulation of β-catenin and c-MYC by GSK-3β inhibition triggered apoptosis in RAS-driven cancer cells and suppressed tumor growth^43^. Likewise, we have shown that MAPK activation induced by FGF2 is also detrimental to KRAS-driven cancer cells and leads to replication stress in an oncogenic KRAS-dependent manner^44^. MAPK pathway activity is suppressed by DUSP-mediated (Dual-Specificity Phosphatases) dephosphorylation^45^. Inhibition of DUSP1/6 induced toxic levels of MAPK activity and triggered cell death in a panel of *Egfr*- and *Kras*-driven lung adenocarcinoma cells^46^. Along the same lines, PP2A phosphatases suppress different oncogenic pathways, and previous data show that inhibition of these phosphatases by LB-100 is toxic to multiple cancer models^14^. Our data confirm the activation of oncogenic signaling and stress-response pathways induced by LB-100. Notably, despite the multitude of cellular processes in which PP2A is implicated^11, 12^, unbiased analyses of our CRISPR screens place WNT/ β-catenin and MAPK signaling at the core of LB-100 toxicity. Because many of the “stress hallmarks” of cancer^8^ were mobilized by LB-100 in CRC cells, the tailored stress-focused drug library was instrumental in identifying CHK1 and WEE1 as LB-100 synthetic lethality targets. We focused on WEE1 in follow up experiments because of the superior clinical development of WEE1 inhibitors compared to those targeting CHK1^21^.

We observed synergy between LB-100 and the WEE1 inhibitor adavosertib across different colon cancer, pancreatic cancer, and cholangiocarcinoma cell models. This suggests there is not a strong context-dependency for this drug combination, facilitating future clinical development. Synergy *in vitro* does not necessarily translate into synergy *in vivo*. Yet, we found higher synergy scores when adavosertib is combined with LB-100 compared to combinations with doxorubicin or gemcitabine across the models tested. This is noteworthy because modest single-drug efficacy has hindered the clinical development of adavosertib and other WEE1 inhibitors so far. Combinations with chemotherapy tend to increase efficacy but also toxicity^47^. In preclinical models, LB-100 improved the efficacy of chemotherapy drugs without compounding the toxicity^14^. This may point towards a better toxicity profile for the LB-100 + adavosertib combination in patients.

The mitotic defects observed are in agreement with previous studies using WEE1 inhibitor combinations. For instance, Aarts and coworkers showed that adavosertib forces S-phase cells into premature mitoses if DNA replication is stalled by gemcitabine^29^. PARP^48^ or ATR^49^ inhibitors also result in replication stress and lead to similar aberrant mitoses in combination with WEE1 inhibition. Mechanistically, these studies have in common the induction of replication stress, priming cancer cells for defective mitoses. In our models, LB-100 also induced replication stress, and the combination with adavosertib led to premature mitoses. Downregulation of DNA repair signaling has been proposed to underlie the replication stress induced by LB-100^50^. Furthermore, PP2A regulates various mitotic proteins, and LB-100 has been shown to induce mitotic catastrophe by deregulating the activity of these proteins^51^. However, the upregulation of oncogenic signaling per se is sufficient to trigger both replication stress^52^ and a myriad of mitotic defects^53, 54^. This notion aligns with our data showing that WNT/β-catenin and MAPK signaling modulate LB-100 toxicity. Identifying the precise mechanistic contribution of each of these effects of LB-100 to the synergy with adavosertib requires further research. Nonetheless, our data show complementary effects of LB-100 and adavosertib on replication stress and mitosis, explaining the efficacy of the combination, and supporting further clinical investigation.

That LB-100 and adavosertib combination showed synergy and no clear context-dependency in the cancer models tested is promising, but also raises questions about efficacy vs. toxicity in vivo. Both drugs have been tested in animal models and clinical trials^14, 25^ but not in this combination. In our study, we found consistent tumor suppression for the combination in clinically relevant CRC and CCA patient-derived tumors, despite different genetic backgrounds, previous treatments, and tissue of origin. Moreover, the fact that the three orthotopic CRC PDXs showed histopathological regression with a reduced proportion of cancer cells in the tumor when treated with the combination is encouraging. The combination was tolerated in immunodeficient mice, however, tolerability in mice does not always translate to humans, and the safety of this combination must be carefully addressed in the clinic.

A potential concern of the approach proposed here is that activation of oncogenic signaling might cause normal or pre-cancerous cells to proliferate in patients. However, normal cells have effective feedback mechanisms to limit oncogenic activity and uncontrolled proliferation. Higher levels of oncogenic signaling and less effective feedbacks to control that signaling are hallmarks of cancer^1^. Consistent with this, mice displaying constitutive ERK1/2 activation due to *Dusp6* knockout do not show increased tumorigenesis^55^. Moreover, results from a phase 1 clinical trial indicate a favorable toxicity profile for LB-100^16^. This scenario suggests a therapeutic window to exploit this hyperactivation, which is supported by our animal data.

Arguably the most appealing aspect of the therapeutic approach described here is that the stress cancer cells need to evade to become resistant is the hyperactivated oncogenic signaling itself. As such, the therapy could select for secondary events that reduce oncogenic signaling and attenuate the malignant phenotype. Indeed, we found downmodulation of oncogenic signaling and reduced proliferation in CRC models after acquiring resistance to the combination. Surprising also was the reduced aneuploidy, a common trait of malignant cells, observed in these combination-resistant cells compared to their parental counterparts. Analyzing the transcriptional “identity” of individual cells after acquired resistance revealed an overall trend towards less malignant phenotypes, despite evident heterogeneity. It is reasonable to assume that these different adaptations identified here (reduced aneuploidy, reduced WNT/β- catenin, MYC, and E2F signaling) cooperate to reduce the tumorigenicity of these resistant cells that we observed *in vivo*. How prevalent this tumor-suppressive acquired resistance will be across different cancer models and drugs activating oncogenic signaling remains to be investigated. Our data are consistent with a model in which further reinforcement of the very oncogenic signaling that underlies the oncogenic phenotype may force the cancer cells to give up signals that fuel oncogenic growth to escape the therapy.

## AUTHOR CONTRIBUTIONS

M.H.D. and R.B. conceived the study. M.H.D., A.F., J.M.F.N., and R.B. wrote the manuscript. M.H.D., A.F., S.W., J.M.F.N., F.v.G., H.J..K., S.M., D.A.V., B.M., A.D., E.v.D., C.P., M.S.S., R.J., A.M.S., E.M., A.V., R.A.M.W., T.E.P.T., J.D.C., J.A.R., and P.S. conceived, performed, and/or analyzed experiments throughout the manuscript. C.S., R.S., and S.Y. established the patient-derived models tumors. C.L. analyzed RNAseq and screens data. S.Mo. analyzed single-cell RNAseq data. M.H.D., W.Q., J.S.K., H.A.A, H.t.R., A.v.O., H.J., Ro.B., A.Vil., R.H.M., and R.B. supervised the study.

## ACKNOWLEDGEMENTS

We thank the members of the Bernards laboratory for helpful discussion and thoughtful feedback. We thank the genomics core facility of the Netherlands Cancer Institute (NKI). This work was supported by an institutional grant of the Dutch Cancer Society and of the Dutch Ministry of Health, Welfare and Sport and by the Oncode Institute, by grant ERC-787925 from the European Research Council (R.B.), by a research grant from Lixte Biotechnology (R.B), by grants 2019/10753-2 and 2020/10277-3 from the São Paulo Research Foundation (M.d.S.), by grant 82222047 from the National Natural Science Foundation of China and 22XD1423100 from Program of Shanghai Academic/Technology Research Leader (H.J.), by grants PI19-01320 and PI22-00548 funded by the Instituto de Salud Carlos III (ISCIII) and co-funded by the European Union, and CERCA Program/Generalitat de Catalunya (2017SGR449 and 2021 SGR 00184) (A.V.), and by grant (2013/07467-1) from the São Paulo State Foundation-FAPESP: CeTICS-Grant (H.A.A.)

## METHODS

### Cell lines and culture

The human cell lines SW-480, HT-29, DLD1, HCT116, LoVo, RKO, AspC-1, MIA-PaCa-2, PANC-1, YAPC, EGI-1, and SSP-25 were obtained from ATCC. DiFi cell line were a gift from Alberto Bardelli. HuCC-T1 and RBE cell lines were provided by Erasmus University. All cell lines were cultured in RPMI medium (except EGI-1, that was cultured in DMEM), supplemented with 10% FBS, 1% penicillin/streptomycin and cultured at 37°C in 5% CO_2_. The cell lines were authenticated by short-tandem-repeat DNA profiling. Mycoplasma tests were performed every 2-3 months. HT-29 and SW-480 resistant cells (HT-29-R and SW-480-R) were established by culturing the parental cells in the presence of increasing concentrations of the combination of LB-100 and adavosertib, up to the maintenance concentrations of 6 µM LB-100 and 600 of nM adavosertib.

### Dose-response assays

All drug-response assays were performed in triplicate, using black-walled 384-well plates (Greiner #781091). Cells were plated with a 20% density (approximately) and incubated overnight for attachment. Drugs were then added to the cells using the Tecan D300e digital dispenser. 10 µM phenylarsine oxide was used as positive control (0% cell viability) and DMSO was used as negative control (100% cell viability). 3-5 days later, resazurin (Sigma #R7017) was added to the plates. After 1-4 hours incubation (depending on the cell line), fluorescence (560_Ex_/590_Em_) was recorded using the EnVision (Perkin Elmer).

### Compounds

LB-100 #206834 and adavosertib (MK-1775) #201912 were purchased from Medkoo Biosciences. GDC-0575 #HY-112167A, doxorubicin #HY-15142, and gemcitabine #HY-17026 were purchased from MedChemExpress.

### IncuCyte-based proliferation and caspase 3/7 activity assays

All IncuCyte assays were performed at least in triplicate, using black-walled 96-well plates (Greiner #655090). Cells were plated at a low density and incubated until attachment. Plates were then placed in the in the IncuCyte ZOOM (Essen Bioscience) which imaged the cells every 4 h. Approximately 12-16 h after plating drugs were added to the cells using the Tecan D300e digital dispenser, as indicated. Phase-contrast images were collected and analyzed to detect cell proliferation based on confluence. When indicated, IncuCyte® Caspase-3/7 green apoptosis assay reagent (Essen Bioscience #4440) was also added to the culture medium. Here, green fluorescent images were also collected and analyzed (by dividing the detected green fluorescence confluence by the phase-contrast confluence) to detect caspase 3/7 activity as a proxy for apoptosis.

### Crystal violet long-term viability assays

Cells were plated at a low density in 12-well plates and incubated overnight for attachment. Drugs were then added to the cells using the Tecan D300e digital dispenser, as indicated. The culture media/drugs were refreshed every 2-3 days. When control wells were confluent, cells were fixed and stained for 20-30 minutes using a 1% formaldehyde (Millipore #104002) /1% methanol (Honeywell #32213)/0.05% crystal violet (Sigma #HT90132) solution in phosphate-buffered saline (PBS). Plates were then washed in water and left to dry before scanning.

### Attached vs anchorage-independent proliferation assays

For HT-29, an equal number of cells were plated in parallel on regular 12-wells plates (TPP #92006) for attached proliferation; and on cell-repellent 12-wells plates (Costar #3471) for anchorage-independent proliferation. The same procedure was used for SW-480 cells but using 96-wells (Greiner #655180) for attached and 96-wells (Greiner #655970) for anchorage-independent proliferation. After 5-6 days, the same initial numbers of cells were seeded in previously empty wells to provide the respective T0 readings. CellTiter-Glo 3D^®^ (Promega #G9682) was then used according to the manufacturer’s protocol to estimate the relative endpoint cell proliferation on both growth conditions. Results are expressed in fold change over T0.

### Synergy matrices

All synergy matrices were performed in triplicate, using black-walled 384-well plates (Greiner #781091). Cells were plated with at 10-20% density and incubated overnight for attachment. Drugs were then added to the cells using the Tecan D300e digital dispenser. 10 µM phenylarsine oxide was used as positive control (0% cell viability) and DMSO was used as negative control (100% cell viability). 5 days later, resazurin (Sigma #R7017) was added to the plates. After 1-4 hours incubation (depending on the cell line), fluorescence (560_Ex_/590_Em_) was recorded using the EnVision (Perkin Elmer). The readings were then normalized using the positive and negative controls to express the relative viabilities. The normalized values for each concentration of single drugs or drug combinations were uploaded on https://synergyfinder.org to calculate the respective synergy scores (ZIP).

### Western blots

After the indicated culture period and drug treatment, cells were washed with cold PBS, then lysed with RIPA buffer (25mM Tris-HCl, pH 7.6, 150 mM NaCl, 1% NP-40, 1% sodium deoxycholate, 0.1% SDS) containing Complete Protease Inhibitor cocktail (Roche) and phosphatase inhibitor cocktails II and III (Sigma). Samples were then centrifuged for 15 min at 15,000 x g at 4°C and supernatant was collected. Protein concentration of the samples was normalized after performing a Bicinchoninic Acid (BCA) assay (Pierce BCA, Thermo Scientific), according to the manufacturer’s instructions. Protein samples (denatured with DTT followed by 5 min heating at 95°C) were then loaded in a 4-12% polyacrylamide gel. Gels were run (SDS-PAGE) for approximately 40 min at 180 volts. Proteins were transferred from the gel to a nitrocellulose membrane at 350 mA for 90-120 min. After the transfer, membranes were incubated in blocking solution (1% BSA/1% non-fat dry milk in TBS with 0.1% Tween-20 (TBS-T). Subsequently, membranes were probed with the primary antibodies in 5% BSA (Bovine Serum Albumin) overnight at 4°C. Membranes were then washed 3 times for 10 min with TBS-T, followed by 1 h incubation at room temperature with the secondary antibody (HRP conjugated) in blocking solution. Membranes were again washed 3 times for 5 min in TBS-T. Finally, a chemiluminescence substrate (ECL, Bio-Rad) was added to the membranes and the signal was imaged using the ChemiDoc-Touch (Bio-Rad).

The following antibodies were purchased from Cell Signaling Technology: Phospho-p44/42 MAPK (Erk1/2) (Thr202/Tyr204) #4377, p44/42 MAPK (Erk1/2) #4695, Phospho-c-Jun (Ser73) #3270, c-Jun #2315, Phospho-4E-BP1 (Thr37/46) #2855, 4E-BP1 #9644, Active β-Catenin (non-phospho) (Ser33/37/Thr41) #8814, β-Catenin #9562, Axin2 #2151, Phospho-Histone H2A.X (Ser139) #2577, Histone-H2A.X #7631, Histone H3 #4499, Phospho-Chk1 (Ser345) #2348, Chk1 #2360, IRE1α (14C10) #3294, HIF-1α #3716, PARP #9542, and c-Myc #5605. The antibodies α-Tubulin #T9026, Vinculin #V9131 were purchased from Sigma Aldrich. Phospho-IRE1 alpha (Ser724) #PA1-16927 was purchased from Thermo Fisher. BCL9L #ab113110 was purchased from Abcam. Phospho-Histone H3 (Ser10) #06-570 was purchased from MERCK (Upstate).

### Stress-focused drug screens

To compose the stress-focused drug library, we considered the following stresses often associated with malignant phenotypes: DNA damage stress, oxidative stress, mitotic stress, proteotoxic stress, metabolic stress, and senescence/apoptosis evasion^8^. Each compound was selected based on published data showing either induction of the respective stress or inhibition of the cellular responses to it. The range of doses was tailored according to published data about the compound when available. On day 0, HT-29 or SW-480 cells were plated in black-walled 384-well plates (Greiner #781091) at 10-20% confluence. On the next day, the plates were divided in control and LB-100 treated arms. LB-100 (2.5 µM) was added to the LB-100 treated arms and then each compound was added on both arms in 15 different doses without replicates using the MICROLAB STAR liquid handling workstation (Hamilton). In each plate, phenylarsine oxide (10 µM) was used as positive control (0% cell viability) and DMSO was used as negative control (100% cell viability). After 3 days, resazurin (Sigma #R7017) was added to the plates. After 1-2 hours incubation, fluorescence (560_Ex_/590_Em_) was recorded using the EnVision (Perkin Elmer). The readings of each plate were then normalized using the positive and negative controls already taking into account any effect of LB-100 alone. The normalized values of each drug/dose were used to build drug-response curves in the absence or presence of LB-100. The area under the curve (AUC) difference for each compound in the presence of LB-100 (2.5 µM) relative to untreated controls are shown.

### RNAseq and gene set enrichment analysis

After plating and attachment overnight, HT-29 and SW-480 cells were treated with LB-100 (2.5 µM) for the indicated time points. For harvesting, the cells were scraped into 15 ml tubes, washed once with cold PBS and homogenized in RLT buffer (Qiagen #79216). Total RNA was isolated using the RNeasy Mini Kit (74106, Qiagen) including an on column DNase digestion (79254, Qiagen), according to the manufacturer’s instructions. Quality and quantity of the total RNA was assessed on the 2100 Bioanalyzer instrument following manufacturer’s instructions “Agilent RNA 6000 Nano” (G2938-90034, Agilent Technologies). Total RNA samples were subjected to TruSeq stranded mRNA library preparation, according to the manufacturer’s instructions (Document # 1000000040498 v00, Illumina). The stranded mRNA libraries were analyzed on a 2100 Bioanalyzer instrument following the manufacturer’s protocol “Agilent DNA 7500 kit” (G2938-90024, Agilent Technologies), diluted to 10nM and pooled equimolar into multiplex sequencing pools for paired end sequencing on the NovaSeq 6000 Illumina sequencing instrument. Paired-end sequencing was performed using 54 cycles for Read 1, 19 cycles for Read i7, 10 cycles for Read i5 and 54 cycles for Read 2, using the NovaSeq6000 SP Reagent Kit v1.5 (100 cycles) (20028401, Illumina).

For the analysis, the RNAseq count data was normalized using a relative total sizefactor. The dataset was then filtered for protein coding genes. Log2 fold change and mean values were calculated for each timepoint of treatment compare to untreated. Before running a specific GSEA analysis, the data was filtered for low counts using a filter of mean greater or equal to 10 in order to obtain robust scores. A GSEA analysis for the Hallmarks geneset and KEGG geneset of the molecular signature database^56^ was performed using the R-package fGSEA^57^ which calculates a NES score and a p-value^56^.

### Single-cell RNAseq

The single-cell RNA-seq data was generated in four runs: HT-29 treatment parental (S1), HT-29 combination-resistant (S2), SW-480 parental (S3), and SW-480 combination-resistant (S4). Parental cells were harvested untreated and resistant cells were harvested 24h after the washout of the drugs. For the single cell 5’ sequencing library preparation, the Chromium Controller platform of 10X Genomics was used for single cell partitioning and barcoding. Each cell’s transcriptome was barcoded during reverse transcription, pooled cDNA was amplified and Single Cell 5’ Gene Expression libraries were prepared according to the manufacturer’s protocol (CG000331 Rev C, 10X Genomics). All libraries were quantified and normalized based on library QC data generated on the Bioanalyzer system according to manufacturer’s protocols (G2938-90321 and G2938-90024, Agilent Technologies). Based on the expected target cells counts, a balanced library pool of all samples was composed. The Single Cell 5’ Gene Expression library pool was quantified by qPCR, according to the KAPA Library Quantification Kit Illumina® Platforms protocol (KR0405, KAPA Biosystems). Paired end sequencing was performed on a NovaSeq 6000 Instrument (Illumina) using a NovaSeq 6000 S1 Reagent Kit v1.5 100 cycles (cat. no. 20028319, Illumina), using 26 cycles for Read 1, 10 cycles for Read i7, 10 cycles for Read i5 and 92 cycles for Read 2. This resulted in an average sequencing depth of 65.000 median reads per cell.

The resulting data was then transformed to FASTQ format and aligned to reference genome (*Homo sapiens* hg38) using the commercial CellRanger 7.0.1 pipeline. For the downstream bioinformatic analyses, we employed the following pipeline per cell line: According to standard QC metrics (RNA content, mitochondrial percentage), cells with low quality were first filtered out, and potential doublets were removed using scDblFinder (v1.10.0)^58^. Dataset were then integrated using Seurat CCA alignment strategy (v4.2.0)^59^, which yielded both UMAP projections^60^ and Leiden clusters^61^. Cluster-specific markers were derived using DESeq2 (v1.36.0)^62^ and cluster-specific enrichment was assessed using the GSEApy (v1.0.1) EnrichR pipeline^63, 64^. Finally, we estimated pathway-activity scores using UCell (v2.0.1)^65^.

### CRISPR screens

#### CRISPR knockout screens

The appropriate number of cells to achieve a 250-fold representation of the Brunello library for all the screen arms and replicates were transduced at approximately 50% confluence in the presence of polybrene (8 μg/ml) with the appropriate volume of the lentiviral-packaged sgRNA library. Cells were incubated overnight, followed by replacement of the lentivirus-containing medium with fresh medium containing puromycin (2 μg/ml). The lentivirus volume to achieve a MOI of 0.3, as well as the puromycin concentration to achieve a complete selection in 3 days was previously determined. After puromycin selection, cells were split into the indicated arms/replicates (for each arm, the appropriate number of cells to keep a 250-fold representation of the library was plated at approximately 10-20% confluence) and a T0 (reference) time point was harvested. Cells were maintained as indicated and, in case a passage was required, cells were reseeded at the appropriate number to keep at least a 500-fold representation of the library. Cells were harvested after about 8 population doublings, washed with PBS, pelleted and stored at -80°C until DNA extraction.

Genomic DNA was extracted (Zymo Research, D3024) from cell pellets according to the manufacturer’s instructions. For every sample, DNA was quantified and the necessary DNA to maintain a 250-fold representation of the library was used for subsequent procedures (for this, we assumed that each cell contains 6.6 pg genomic DNA). Each sample was divided over 50 μl PCR reactions (using a maximum of 1 µg genomic DNA per reaction) using barcoded forward primers to be able to deconvolute multiplexed samples after next generation sequencing. PCR mixture per reaction: 10 μl 5x HF Buffer, 1 μl 10 μM forward primer, 1 μl 10 μM reverse primer, 0.5 μl Phusion polymerase (ThermoFisher, F-530XL), 1 μl 10mM dNTPs, adding H2O and template to 50 μl. Cycling conditions: 30 sec at 98°C, 20× (30 sec at 98°C, 30 sec at 60°C, 1 min at 72°C), 5 min at 72 °C. The products of all reactions from the same sample were pooled and 2 μl of this pool was used in a subsequent PCR reaction using primers containing adapters for next generation sequencing. The same cycling protocol was used, however this time for 15 cycles. Next, PCR products were purified using the Bioline ISOLATE II PCR and Gel Kit (GC Biotech, BIO-52060) according to the manufacturer’s instructions. DNA concentrations were measured and, based on this, samples were pooled equimolarly.

The pool of amplified sgRNA sequences was sequenced on an Illumina HiSeq 2500 with high-output kit (Single-Read, 65 bp). The reads were mapped to the unique barcodes used for each sample and sgRNA sequences of the Brunello library. Mapped read-counts were subsequently used as input for the further analyses.

#### CRISPRa screen

HT29 dCas9-VP64 cells were generated by lentiviral transduction of HT29 cells with Lenti-dCas9-VP64-Blast (Addgene, 61425) in the presence of 8 μg/ml polybrene (Santa Cruz, sc-134220A) and subsequent selection with 10 µg/ml blasticidin (InvivoGen, ant-bl). Clonal derivatives of the HT-29 dCas9-VP64 cell line were established by limited dilution.

HT29 dCas9-VP64 clone E cells were transduced with lentivirus of Calabrese pooled human CRISPRa library set A and B (Addgene, 92379 and 92380) separately, in the presence of 8 μg/ml polybrene (Santa Cruz, sc-134220A), and at a multiplicity of infection of approximately 0.3. Following 3 days of selection with 2 µg/ml puromycin (Gibco, A11138-03), reference samples were collected (t=0) and cells were separated into different treatment arms. Cells were subsequently cultured in the presence of 5 µM LB-100 (2 and 3 replicates for set A and B, respectively) or DMSO (vehicle control; 3 replicates for each set) while maintaining at least 16 million cells per replicate at all times, ensuring a 250-fold representation of each library set. After both arms reached at least 12 population doublings, cells were collected and stored as pellet at -80°C.

Genomic DNA was isolated using the Gentra® Puregene® kit (Qiagen, 158767) following the manufacture’s protocol specified for cultured cells, and dissolved in the hydration solution overnight while shaking at room temperature. DNA yield ranged from 216 µg to 486 µg per sample. The genomic DNA was divided into multiple reactions per sample (50 µg each, using all material) and fragmented at 37°C overnight, using 100 U Ndel enzyme (R0111L) and 50 µl 10X cutSmart® buffer (B7204S) from New England Biolabs, supplemented to 500 µl with nuclease-free water (ThermoFisher, AM9932). The reactions were heated to 100°C for 10 minutes, and following addition of 500 µl 2 M NaCl, reheated to 100°C for 5 minutes and then immediately snap-frozen in liquid nitrogen. Per tube and prior to thawing, 1 µl of each 10 µM 5’ biotinylated capture oligo (TGCTTACCGTAACTTGAAAGTATTTCGATTTCTTGGCTTTATATATCTTG and TGCTCTCGTGGAGAGGAGCGACGCCATATCGTCTGCTCCCTCGTATTCGC) was added on top of the frozen solution, which was then immediately transferred to a thermoshaker for overnight hybridization at 60°C. To capture hybridized DNA encoding sgRNA sequences, 20 µl Streptavidin T1 Dynabeads (ThermoFisher, 65602) were washed three times with 500 µl wash buffer (1 M NaCl, 10 mM Tris-HCl, pH 8), added to each tube, and incubated under rotation at room temperature for 2 hours. The beads were washed twice with wash buffer and twice with 10 mM Tris-HCl (pH 8). Non-hybridized biotinylated oligonucleotides were digested in 50 µl reactions composed of 44 µl 10 mM Tris-HCl (pH 8), 5 µl 10X Exonuclease buffer, and 1 µl Exonuclease I (New England Biolabs, M0293L), at 37°C for 1 hour. Beads were washed 3 times with 10 mM Tris-HCl (pH 8) and resuspended in 20 µl 10 mM Tris-HCl (pH 8).

Two rounds of PCR were performed to amplify the sgRNA sequences. In the first PCR, distinct forward primers that each encode a unique barcode sequence and facilitate deconvolution of sequence reads of pooled samples (ACACTCTTTCCCTACACGACGCTCTTCCGATCTNNNNNNGGCTTTATATATCTTGTGGAAAGGACG with NNNNNN representing barcode sequences CGTGAT, ACATCG, GCCTAA, TGGTCA, AAGCTA, GTAGCC and TACAAG) were used in combination with a common reverse primer (GTGACTGGAGTTCAGACGTGTGCTCTTCCGATCTCGACGCCATATCGTCTGCT). PCR mixture: 1 μL 10 μM forward primer, 1 μL 10 μM reverse primer, 1 μL 10 mM dNTPs (ThermoFisher, R0193), 0.5 μL Phusion polymerase and 10 μL 5X HF buffer (New England Biolabs, M0530L), supplemented with nuclease-free water to a total volume of 50 μL. PCR cycling conditions: 3 minutes @ 98°C, 20 times (30 seconds @ 98°C, 30 seconds @ 60°C, 30 seconds @ 72°C), and 5 minutes @ 72°C. Per sample, products of individual reactions were pooled and 2 μL of each pool was used as template in the second PCR with conditions similar to the first, but having 15 instead of 20 cycles, to add the p5 and p7 adapter sequences as well as unique indices to discriminate samples of the Calabrese library set A and B (primers: AATGATACGGCGACCACCGAGATCTACACTCTTTCCCTACACGACGCTCTTCCG ATCT and CAAGCAGAAGACGGCATACGAGATNNNNNNGTGACTGGAGTTCAGACGTGTGC TCTTCCGATCT with NNNNNN representing index sequence ACATCG and GCCTAA). The PCR products were purified using the Bioline ISOLATE II PCR and Gel kit (GC Biotech, BIO-52060) following the manufacture’s protocol and pooled by combining 150 ng of each sample.

The pool of amplified sgRNA sequences was sequenced on an Illumina NextSeq with high-output kit (Single-Read, 75 bp). The reads were mapped to the unique barcode and index combination used for each sample and sgRNA sequences of both Calabrese library sets. Mapped read-counts were subsequently used as input for the further analyses.

#### Bioinformatics Analysis

For both type of CRISPR screen, the sequence count data was normalized using a relative total size factor. Statistical comparisons of the conditions treated vs untreated were performed using drugZ^66^. Log2 fold change were calculated based on the median of each of the two conditions. The first criterion for hit selection was a drugZ FDR smaller or equal to 0.25 in treated/untreated comparison. In addition, for negative selection log2 fold change of treated/untreated should be smaller or equal to -1. For positive selection those should be greater or equal to 1.

### String network analysis

The full list of hits from both CRISPR screens showed in figure S2 was inputted on the STRING web tool (https://string-db.org) and analyzed using the default settings. The top 5 Gene Ontology (GO) Biological Processes and Molecular Functions with their respective FDRs are shown.

### Time-Lapse Microscopy

Cells were plated on 8-well glass-bottom dishes (LabTek) and incubated overnight for attachment. Drugs were then added as described and the cells were imaged using a Deltavision deconvolution microscope equipped with a heat chamber. For DNA visualization, cells stably expressed H2B-GFP (obtained by retroviral infection). Images were acquired every 5 min using a 203 (0.25 NA) objective. Z-stacks were acquired with 2-mm intervals. Images were analyzed and processed using Softworx and ImageJ.

### Immunofluorescence staining

Cells were plated on 12mm glass coverslips and incubated overnight for attachment. After the indicated treatments, the cells were fixed for 15 minutes at room temperature in 4% formaldehyde with 0.5% Triton X-100. The mouse anti-alpha-tubulin (Sigma, #t5168) was incubated over night at 4C. Secondary antibodies (Molecular probes, Invitrogen) and DAPI (1 μg/ml) were incubated for 2 hours at room temperature. Coverslips were mounted using ProLong Gold (Invitrogen). Images were taken on a THUNDER Imager 3D Cell Culture van Leica 63× oil lens: Obj. HC PL APO 63×/1.40– 0.60 OIL 11506349.

### Chromosome spreads

Chromosome spreads were prepared from HT-29 cells treated with inhibitors for 16 hours. After that, cells were treated with Nocodazole for 3 hours and harvested by mitotic shake-off. Cells were then incubated with 0.075M of KCl at 37°C for 10 minutes and a drop of fixative (methanol:acetic acid, in a 3:1 ratio made fresh) was added followed by centrifugtion at 1500 rpm for 5 minutes. The supernatants were discarded and the cells were fixed with 1 ml of fixative for 30 minutes, followed by fixative + Dapi (1 μg/ml). The cell suspensions were then dropped from 5 cm distance onto an ethanol cleaned coverslips, dried at room temperature and the chromosome spreads were mounted with ProLong Gold (Invitrogen). Images were acquired using a Thunder deconvolution microscope (Applied Precision) with a 60x 1.40 NA oil objective. Softworx (Applied Precision), ImageJ, Adobe Photoshop and Illustrator CS6 were used to process acquired images.

### Detection of BrdU foci under native DNA conditions

For detection of long fragments of single-stranded DNA (ssDNA), typical of replication stress, BrdU (50 mM) was incorporated into the DNA of exponentially growing cells (HT-29 and SW-480) for 48 h. After that, we washed the coverslips and added fresh media adding the drugs as indicated for 8 h. Next, cells were fixed using 4% of paraformaldehyde in PBS and permeabilized with 0.2% Triton-X 100. BrdU was detected (when accessible) using a purified mouse Anti-BrdU (BD Biosciences) followed by a secondary antibody goat anti-mouse conjugated to Alexa Fluor 488 (Thermo Scientific). To ensure that all cells incorporated BrdU, one additional coverslip for each condition analyzed was prepared to be subjected to DNA denaturation using 2 M HCl (for 15 min), followed by a neutralization step with 0.1 M Borate buffer (100 mM H_3_BO_3_, 75 mM NaCl, 25 mM Na_2_B_4_O_7_·10H2O, pH = 7.4) for 10 min (not shown). Stained coverslips were mounted with VECTASHIELD Mounting Medium with DAPI (Vector Labs). Images were captured using Olympus BX51 fluorescence microscope coupled with a digital camera (XM10; Olympus) and analyzed using OLYMPUS CELL F software (version 5.1.2640, Tokyo, Japan). At least 100 cells were analyzed per coverslip.

### DNA fiber assays

For the DNA fiber assays, after the indicated treatments, cells were labelled with CldU (25 µM, 20 minutes) and IdU (250 µM, 20 minutes). Labelled cells were lysed (200 mM Tris-HCl ph 7.4, 50 mM EDTA, 0.5% SDS), spread onto IHC Microscopy slides (Dako) and fixed for 10 minutes in methanol: acetic acid (3:1). Next, slides were incubated in HCl (2.5 M) for 1 hour and 15 minutes, washed with PBS and incubated in blocking solution (PBS + 1% BSA and 0.1% Tween 20) for 1 hour. Primary rat-anti-BrdU BU1/75 (1:500, Abcam) and mouse-anti-BrdU antibody Clone B44 (1:750, BD Biosciences) were incubated for 1 hour in blocking solution. After washing with PBS, primary antibodies were fixed for 10 minutes using 4% paraformaldehyde. Secondary antibodies (goat-anti-mouse Alexa Fluor 488 and goat-anti-rat Alexa Fluor 555 (both 1:500, Invitrogen) were incubated for 1.5h in blocking solution. Finally, Menzel-Gläser coverslips were mounted onto the slides using Vectashield and imaged using Zeiss AxioObserver Z1 inverted microscope using a Hamamatsu ORCA AG Black and White CCD camera.

### Cell cycle and p-H3 flow cytometry

For bromodeoxyuridine (BrdU)/propidium iodide cell cycle analyses, after the indicated treatments, cells were harvested by trypsinization, washed with cold PBS, and then and fixed in ice-cold 75% ethanol in PBS overnight at 4 °C. BrdU (10 μM) was added 1 hour before harvesting. Fixed cells were washed with PBS and treated with 5M HCl and 0.5% Triton-X100 for 20 min and then washed with 10 mL TrisHCl (pH 7.5). Next, cells were incubated with mouse anti-BrdU (DAKO clone BU20A 1:40) for 1h. After washing with PBS, cells were incubated with polyclonal goat anti-mouse FITC (DAKO F0479 1:20). Finally, cells were washed and resuspended in PBS with propidium iodide (PI) (20 µg/ml) and RNase A (200 µg/ml), incubated at 37 degree for 30 minutes and finally measured on the flow cytometer.

For phospho-histone H3 (S10) staining cells were fixed as described above, washed in PBS, and incubated for 1 h with the conjugated histone antibody (histone H3 S10 Millipore 06-570-AF488). Cells were then resuspended in a propidium Iodide (PI) (50 µg/ml) + RNase A (10 µg/ml) solution in PBS for at least 20 min before analysis in the flow cytometer. For all flow cytometer experiments, data were acquired with Attune NxT flow cytometer (Life Technologies, Carlsbad, CA, USA) and analyzed with flowjo V.10 software (Treestar, Inc., Ashland, OR, USA). At least 20 000 cells per sample were analyzed.

### Animal models

#### Generation of Patient-Derived Orthotopic Xenografts from colorectal tumors and drug treatments

Primary tumors were obtained from Bellvitge Hospital (HUB) and the Catalan Institute of Oncology (ICO) with approval from the Ethical Committee (CEIC Bellvitge Hospital), ethical and legal protection guidelines for human subjects, including informed consent. The experimental design was approved by the IDIBELL animal facility committee (AAALAC Unit1155) under approved procedure 9111. All animal experiments were performed following the guidelines for Ethical Conduct in the Care and Use of Animals as stated in The International Guiding Principles for Biomedical Research Involving Animals, developed by the Council for International Organizations of Medical Sciences.

To establish the orthotopic CRC models from refractory metastatic CRC patients, a small fragment of lung metastases (PDOX1 and PDOX2) or a peritoneal implant (PDOX3) from three different colorectal cancer patients previously treated with fluoropyrimidines-based chemotherapy (see below) were obtained. Briefly, a small tumor piece of 2–4 mm^3^ maintaining tridimensional structure was anchored with Prolene 7.0 to the serosa of the caecum of two five to six-week-old male athymic nude mice (strain Hsd:Athymic Nude-Foxn1nu) purchased from Envigo. After implantation, mice were inspected twice a week. At euthanasia, the tumors were harvested, cut into small fragments, and serially transplanted into new animals for tumor perpetuation or experimental treatment procedures.

PDOX 1 was generated from a lung metastasis of a male patient initially diagnosed with stage III colon adenocarcinoma (MSS, RAS, and BRAF WT). This patient received adjuvant Folfox (first line) and Folfiri + Cetuximab (second line) upon liver relapse; A subsequent relapse in the liver was surgically removed. After subsequent liver and lung (from which the PDOX was generated) progression, Folfox + Cetuximab (third line) were given, obtaining a partial response.

PDOX 2 was generated from a lung metastasis of a male patient initially diagnosed with stage III left colon adenocarcinoma (MSS, RAS, and BRAF WT). This patient received adjuvant capecitabine (first line) and Folfox + panitumumab (second line) upon liver relapse. A subsequent relapse in the lung (from which the PDOX was generated) and adrenal gland were surgically removed. After a liver relapse, Folfiri + aflibercept (third line) were given as neoadjuvant therapy.

PDOX 3 was generated from a male patient initially diagnosed with stage IV colon adenocarcinoma (RAS WT, BRAF V600E mutant). This patient received Folfox as the first-line treatment (partial response) and Cetuximab + Encorafenib + Binimetinib as the second-line (partial response). Upon relapse, a peritoneal implant sample was obtained to generate the PDX, and Folfiri + aflibercept was given as a third-line treatment (partial response).

For the treatment experiments, fragments of PDOX1 (n=25 mice), PDOX2 (n=17 mice), and PDOX3 (n=25 mice) tumors were transplanted into the cecum of mice. When tumors reached a homogeneous palpable size (3 to 5 weeks), mice were randomly allocated into the treatment groups: Vehicles; LB-100 (1.5 mg/kg); Adavosertib (80 mg/kg); and LB-100 + Adavosertib at the same doses. LB-100 was administered by intraperitoneal injection (i.p) on days 1, 3, and 5, while Adavosertib was administered by oral gavage (o.g) on days 1-5 in 7-day cycles. For combined treatments, adavosertib was administered 2-3 hours after LB-100. Drugs were prepared fresh before each daily treatment. Adavosertib was formulated in 2% DMSO+30% PEG 300+5% Tween 80+ddH2O, and LB-100 was dissolved in water. 4h after the last treatment, mice were sacrificed, and tumors were collected, measured, and imaged. Tumour volumes based on calliper measurements were calculated using the modified ellipsoidal formula: tumour volume = ½ length × width^2^. Representative tumor fragments were either frozen in nitrogen or fixed and then processed for paraffin embedding.

#### Cholangiocarcinoma patient-derived xenografts

In compliance with the protocol approved by the Institutional Review Board of Naval Military Medical University Affiliated Eastern Hepatobiliary Hospital and with the informed consent of the participant, fragments of surgically resected tumour tissues from a patient with ICC was used for xenotransplantation (PDX1 = CH-17-0005 FP6). In brief, patient samples were collected, trimmed, cut into 20–30-mm^3^ fragments and implanted subcutaneously in the fore flanks of anaesthetized 6–8-week-old male BALB/c nude mice within 3 h.

Tumour volumes based on caliper measurements were calculated using the modified ellipsoidal formula: tumour volume = ½ length × width^2^. After the tumour volumes reached around 50-100 mm^3^, mice were randomized into the indicated treatment groups. LB-100 (intraperitoneal injection) was given on day 1, adavosertib (oral gavage) was given on days 1-3, in 4-days cycles. All procedures and protocols were approved by the Institutional Animal Care and Use Committee of Shanghai (IACUC NO. 2022-0025).

#### Engraftment of parental vs. combination-resistant cells

The experiment was approved by the Animal Ethics Committee of the Netherlands Cancer Institute. SW-480 parental or SW-480 combination-resistant cells were resuspended in PBS and mixed 1:1 with matrigel (Corning 354230). Three million cells per mice (n=10 per group) were injected subcutaneously into the posterior right flanks of 7-week-old immunodeficient NMRI nude mice. Tumour size was measured 3-times a week by caliper and the volume was calculated by the modified ellipsoidal formula (tumour volume = 1/2(length × width^2^)). Mice were sacrificed at the ethical endpoint of 1500 mm^3^.

### Statistics and reproducibility

With the exceptions of CRISPR screens, drug screens, RNAseq, and single-cell RNAseq, each *in vitro* experiment has been independently reproduced with similar results. GraphPad Prism was used for the statistical analyses.

## SUPPLEMENTARY FIGURE LEGENDS

**Figure S1:**
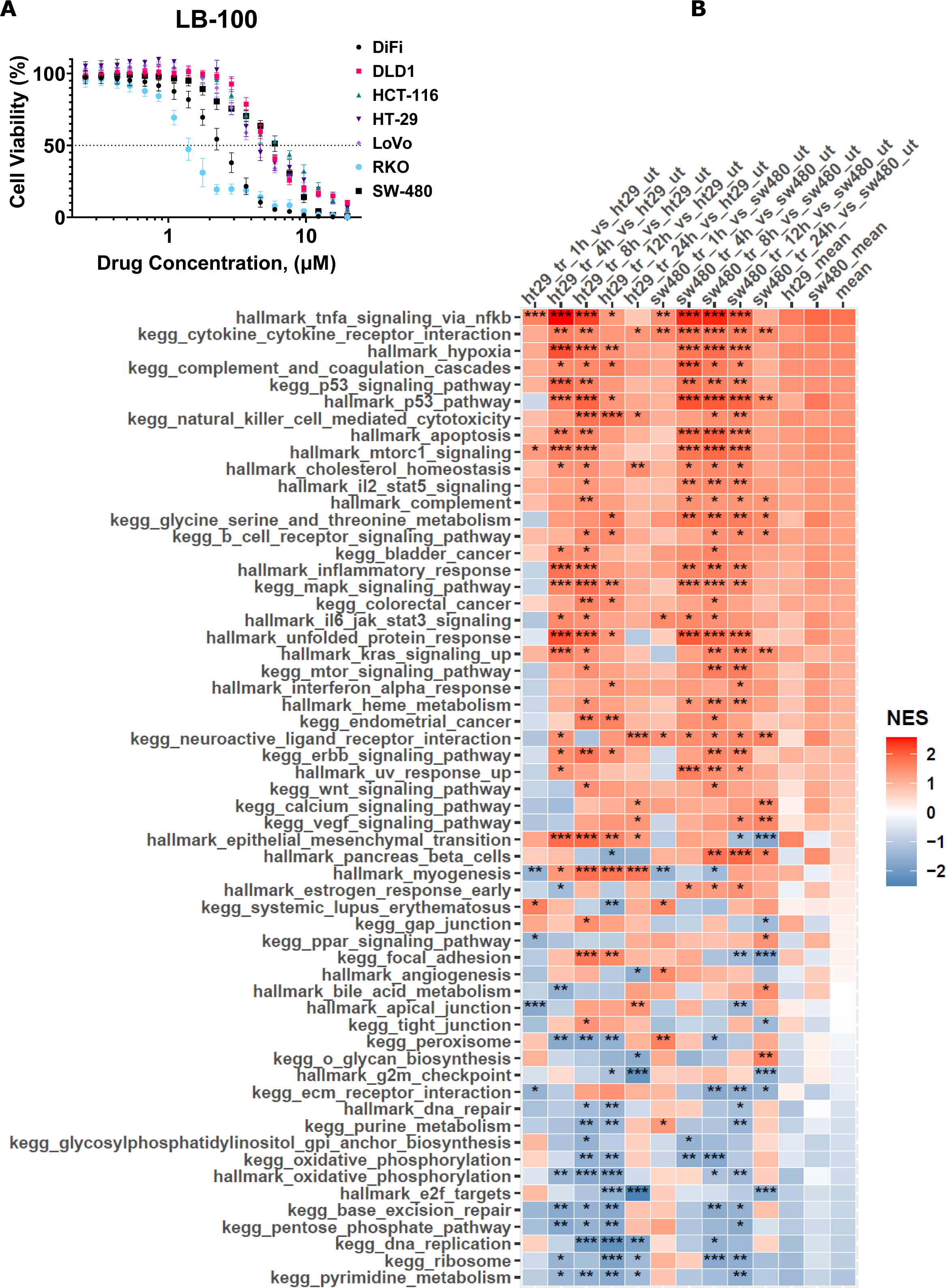
LB-100 engages stress-related, inflammatory response, and mitogenic signaling transcriptional programs in CRC cells. (A) Dose-response assays show the effect of LB-100 in 7 CRC models. Cell viability was estimated by resazurin fluorescence after 5 days in the presence of the drug or DMSO control. The normalized values are plotted. **(B)** The heat map shows all “Hallmarks” and “KEGG” molecular signatures significantly enriched by LB-100 (4 µM) in both HT-29 and SW-480 cells in at least one of the addressed time points. Asterisks indicate significance level (* p-value <0.05, ** p-value <0.01, *** p-value <0.001)

**Figure S2:**
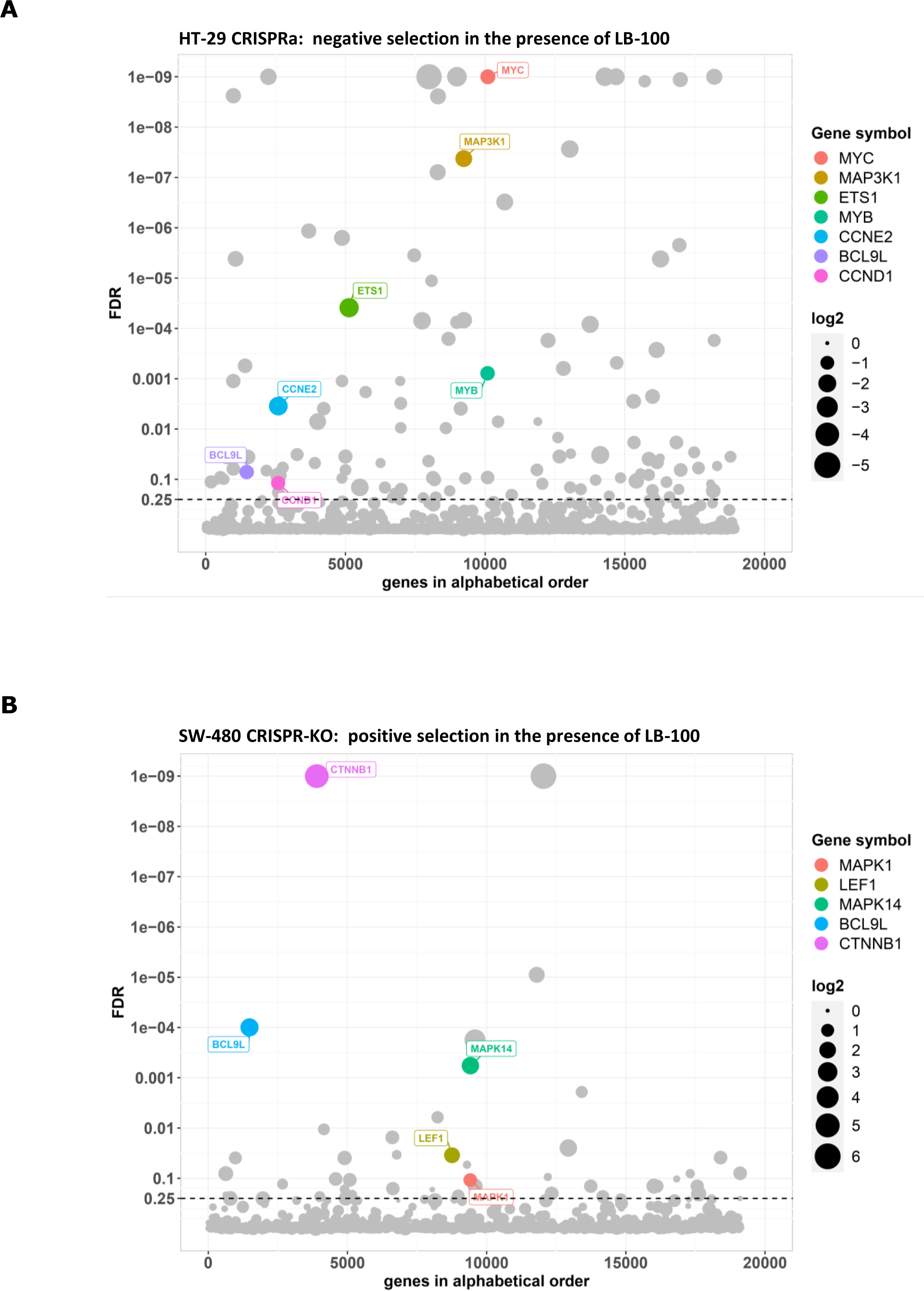
Genome-wide CRISPR screens identify genes modulating LB-100 toxicity in CRC cells. (A) The bubble plot shows genes whose overexpression in the CRISPRa screen was selectively toxic for LB-100-treated (5 µM) cells comparing to untreated controls. 6 different gRNAs per gene were tested in 3 independent replicates. Cells on both conditions were grown for 12 population doublings before DNA harvesting and sequencing. Hits were called based on 0.25 false discovery rate (FDR) and at least 1 log2 fold-change difference between treated and untreated samples. FDR values < 10e-10 are truncated to 10e-10. Only the hits mentioned on the main text are named and colored, the full list of hits is presented on the Supplementary table S2. **(B)** The bubble plot shows genes whose knockout in the CRISPR-KO screen was enriched in LB-100 (6 µM) treated cells comparing to untreated (DMSO) controls. 4 different gRNAs per gene were tested in 3 independent replicates. Cells on both conditions were grown for at least 6 population doublings before DNA harvesting and sequencing. Hits were called based on 0.25 false discovery rate (FDR) and at least 1 log2 fold-change difference between treated and untreated samples. FDR values < 10e-10 are truncated to 10e-10. Only the hits mentioned on the main text are named and colored, the full list of hits is presented on the Supplementary table S3.

**Figure S3:**
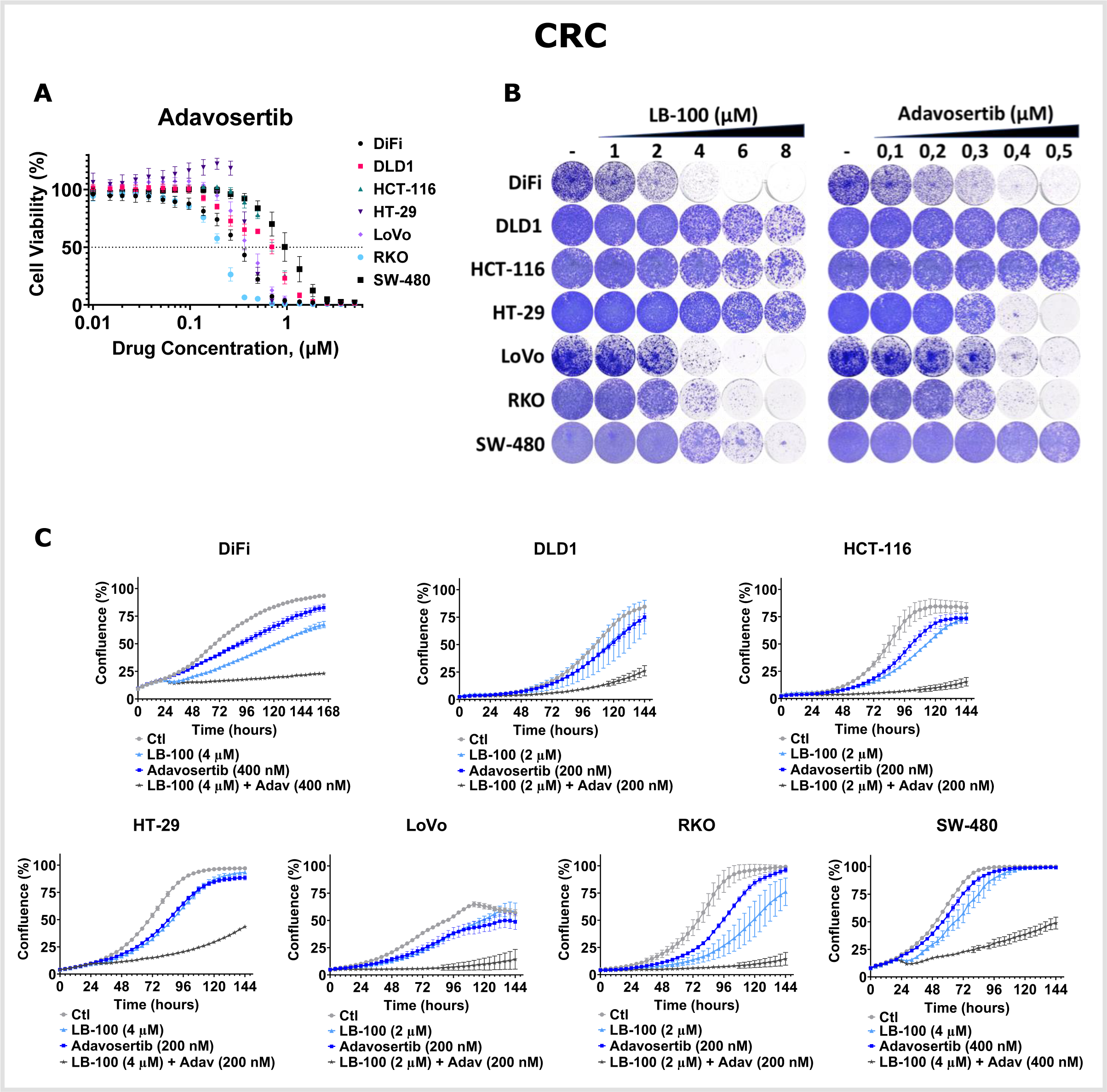
Combined toxicity of LB-100 and Adavosertib in CRC models. (A) Dose-response assays show the effect of Adavosertib in 7 CRC models. Cell viability was estimated by resazurin fluorescence after 5 days in the presence of the drug or DMSO control. The normalized values are plotted. **(B)** Long-term viability assays show 7 CRC models treated with LB-100 or Adavosertib at the indicated concentrations. Treatments were refreshed every 2-3 days, and the cells were grown for 10-14 days before fixing, staining, and imaging. **(C)** IncuCyte-based proliferation assays from 7 CRC models in the absence or presence of LB-100, Adavosertib, or the combination at the indicated concentrations.

**Figure S4:**
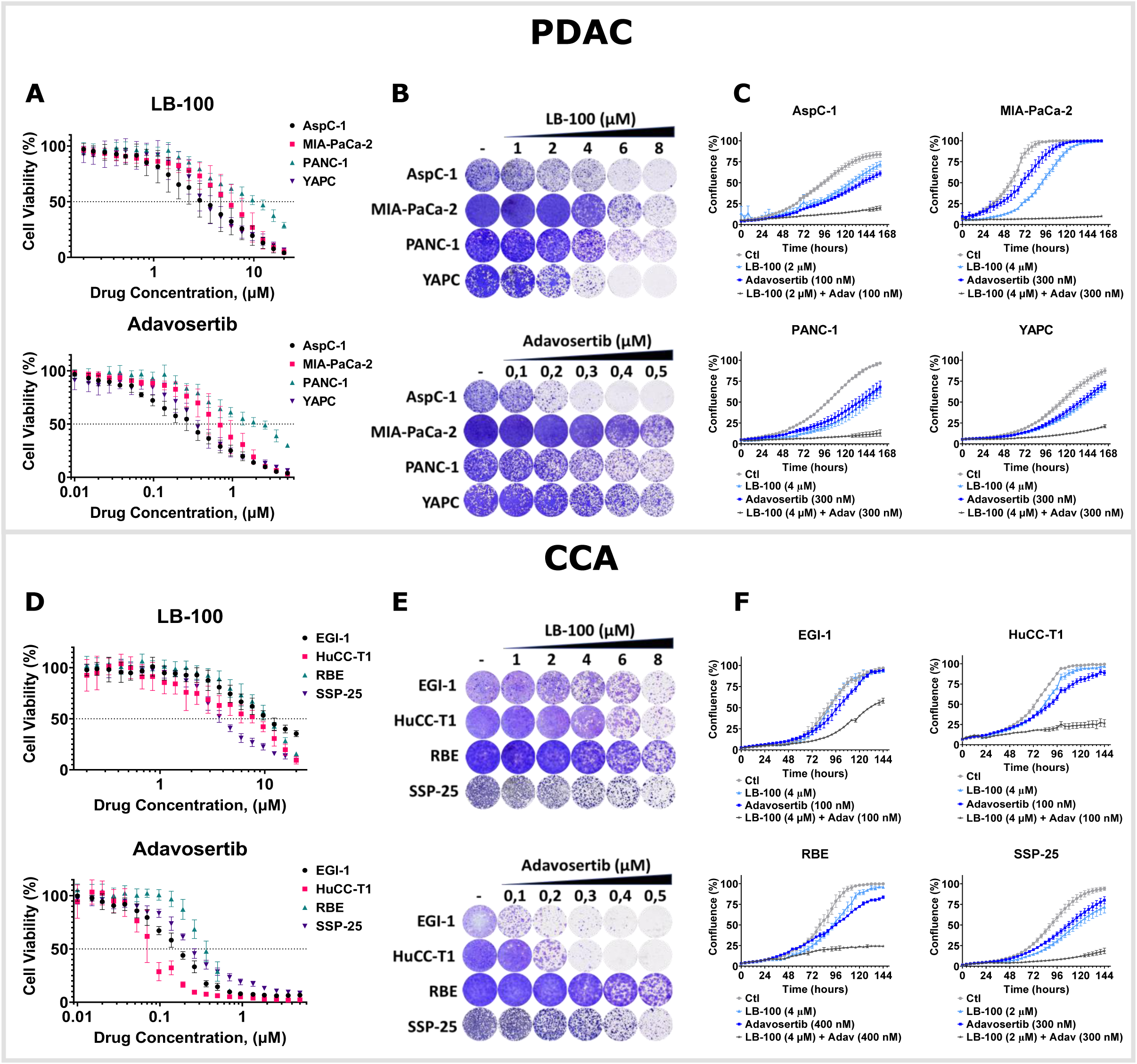
Combined toxicity of LB-100 and Adavosertib in PDAC and CCA models. (A) and **(D)** Dose-response assays show the effect of LB-100 or Adavosertib in 4 PDAC and 4 CCA models, respectively. Cell viability was estimated by resazurin fluorescence after 5 days in the presence of the drug or DMSO control. The normalized values are plotted. **(B)** and **(E)** Long-term viability assays show 4 PDAC and 4 CCA models, respectively, treated with LB-100 or Adavosertib at the indicated concentrations. Treatments were refreshed every 2-3 days, and the cells were grown for 10-14 days before fixing, staining, and imaging. **(C)** and **(F)** IncuCyte-based proliferation assays from 4 PDAC and 4 CCA models, respectively, in the absence or presence of LB-100, Adavosertib, or the combination at the indicated concentrations.

**Figure S5:**
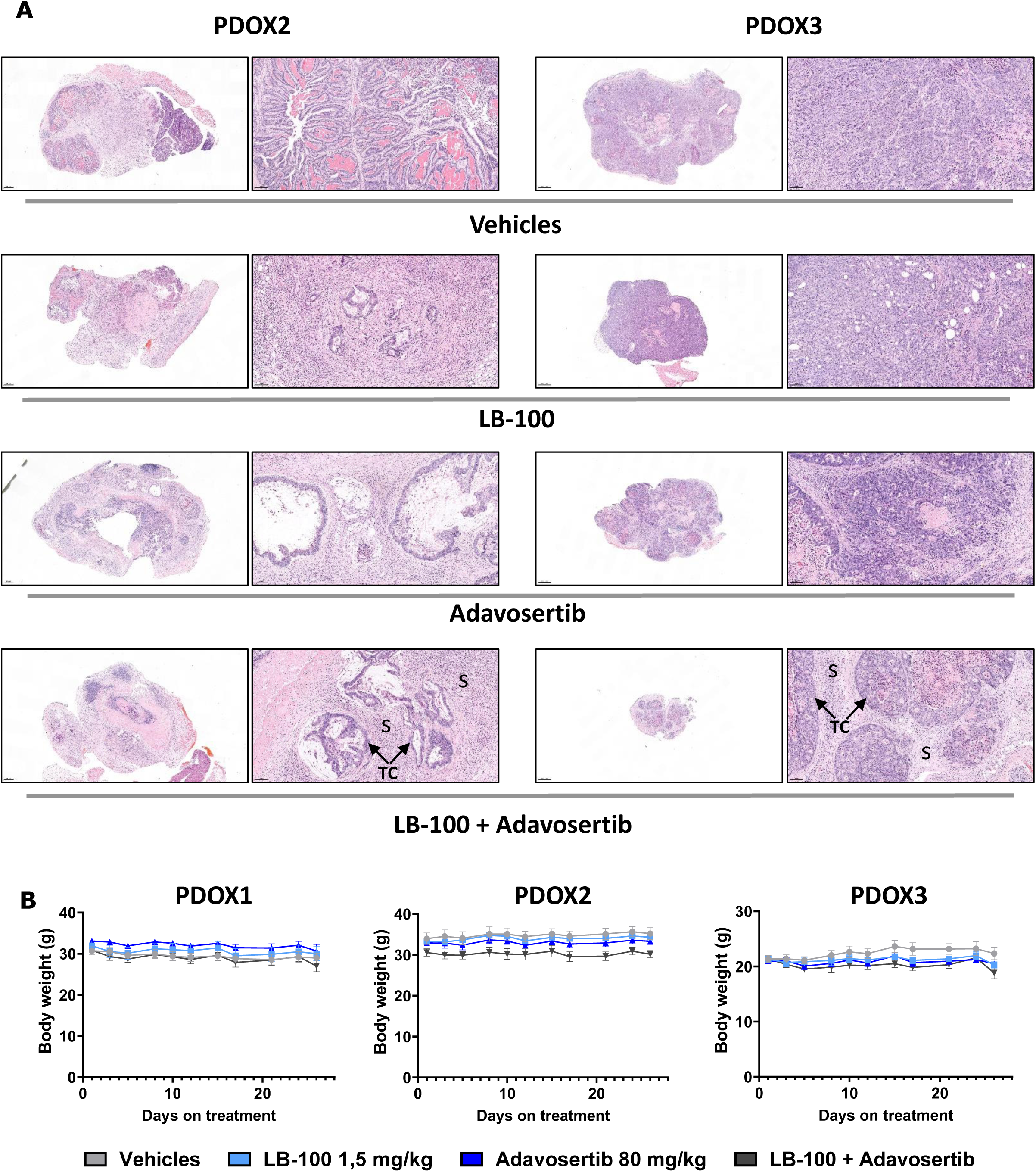
LB-100 and adavosertib induce histological response in orthotopic CRC PDXs. (A) Representative Hematoxylin & Eosin (H&E) stainings at endpoint from PDOX2 and PDOX3 treated as indicated. Original magnification middle images: 15x, scale bar 1000 µm; right images: 200x, scale bar 100 µm. S indicates stroma, and the arrows point to the tumor-cell component. **(B)** Mice body weight variation of the 3 CRC PDXs across the experiments.

**Figure S6:**
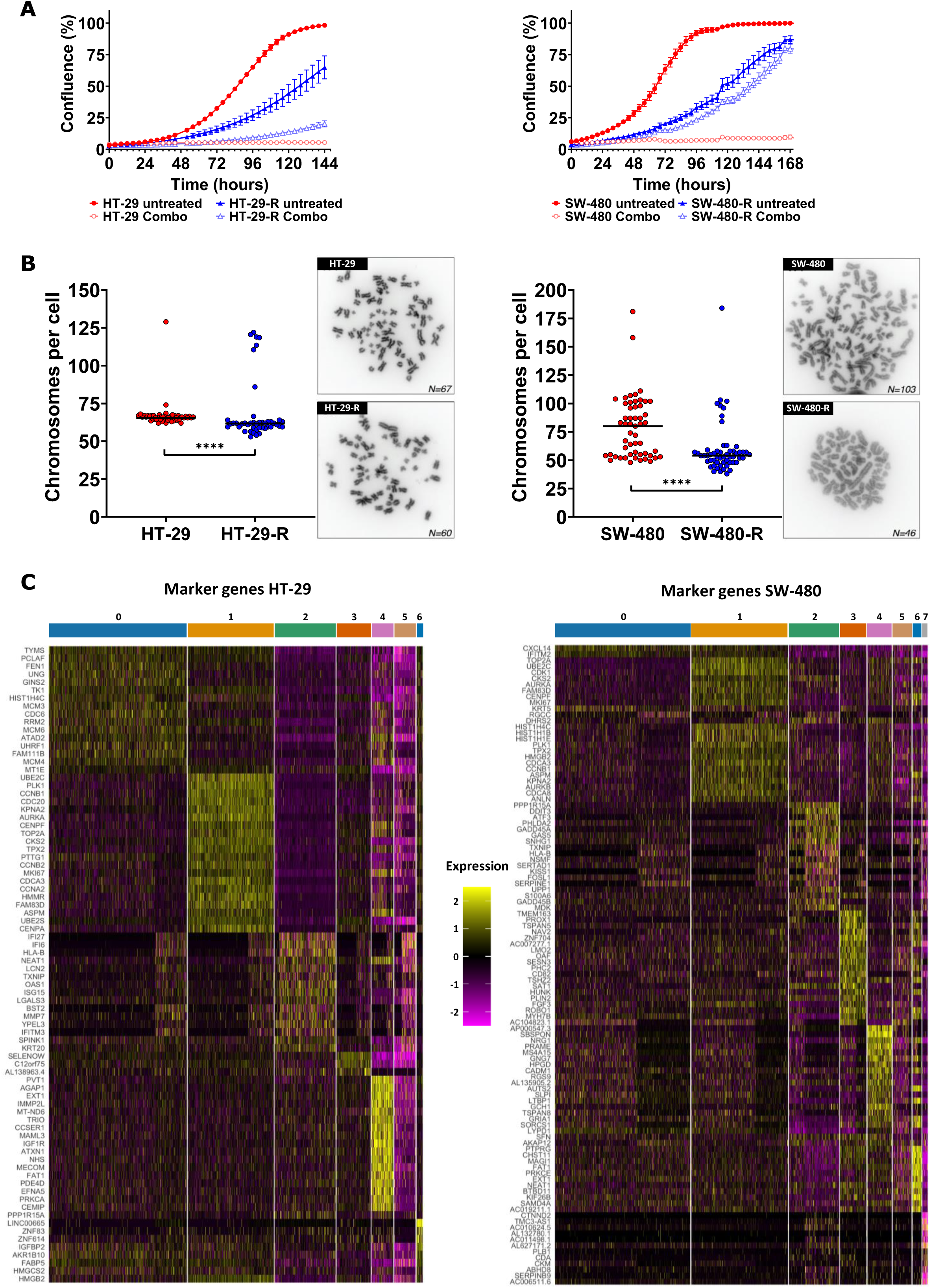
Acquired resistance to the combination of LB-100 and adavosertib suppressed malignant traits in CRC models. (A) IncuCyte-based proliferation assays from HT-29 and SW-480 parental and resistant cells in the absence or presence of the combination (LB-100 4 µM + adavosertib 400 nM). **(B)** Chromosome counting and representative chromosome spreads from HT-29 and SW-480 parental and resistant cells. Nocodazole was added for 3h to block cells in mitosis. Cells were harvested by mitotic shake-off for spreading. Over 40 (HT-29 and HT-29-R) or 50 (SW-480 and SW-480-R) spreads were counted per cell line. Asterisks indicate significance level (**** p-value <0.0001) by two-tailed Mann-Whitney test. **(C)** Heatmaps showing the marker genes of each cluster from the scRNAseq analyses of HT-29 and SW-480 cells.

**Figure S7:**
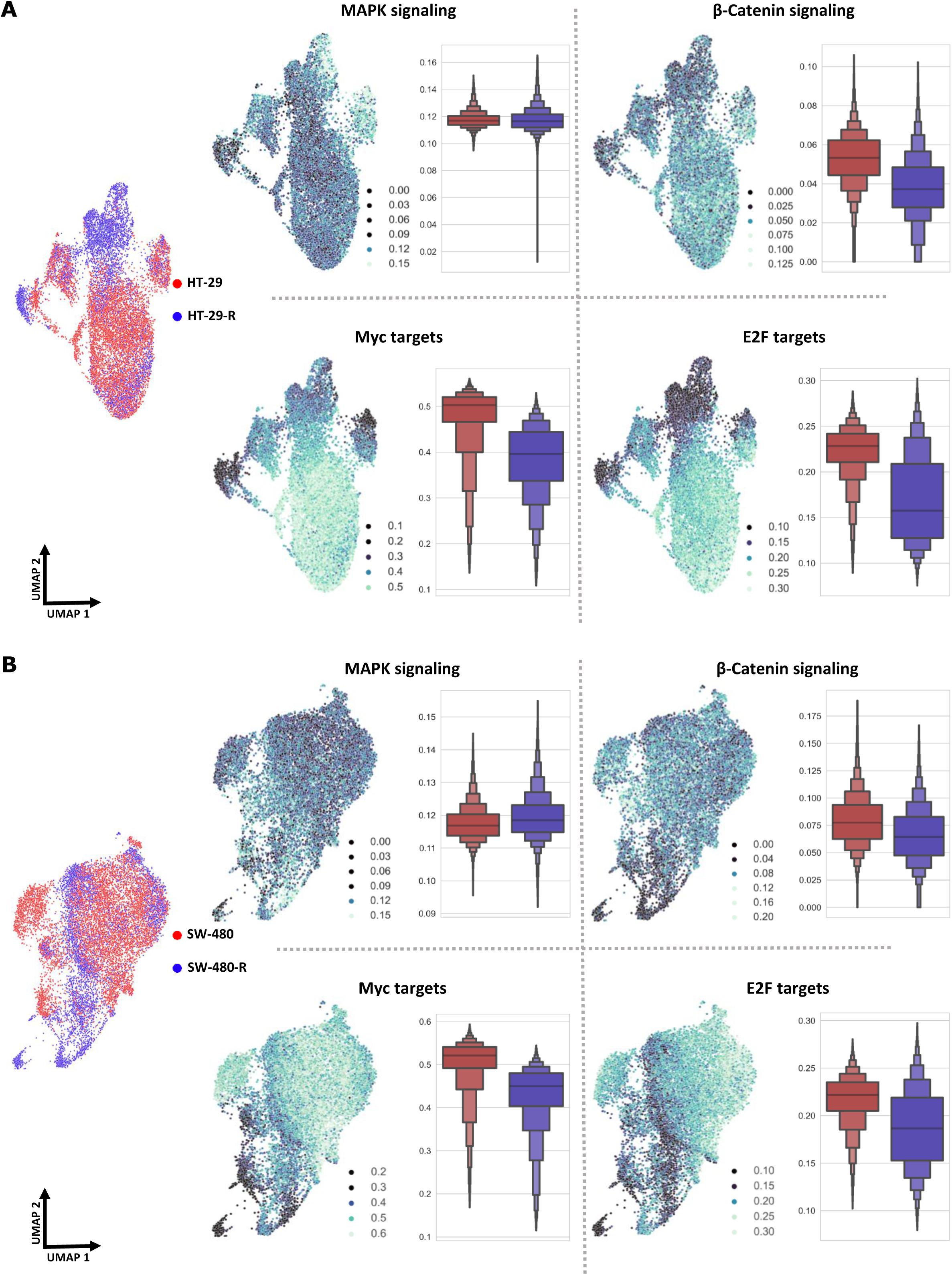
Single-cell RNAseq identify transcriptional signatures downregulated in CRC cells after acquired resistance to the combination of LB-100 and adavosertib. UMAP representations of HT-29 **(A)** and SW-480 **(B)** cells colored by the activity scores for the indicated pathways. UMAPs colored by sample of origin from both cell lines are present of the left for reference. The boxen plots show the pathway activity scores for parental (red) and resistant (blue) cells.

**Figure S8:**
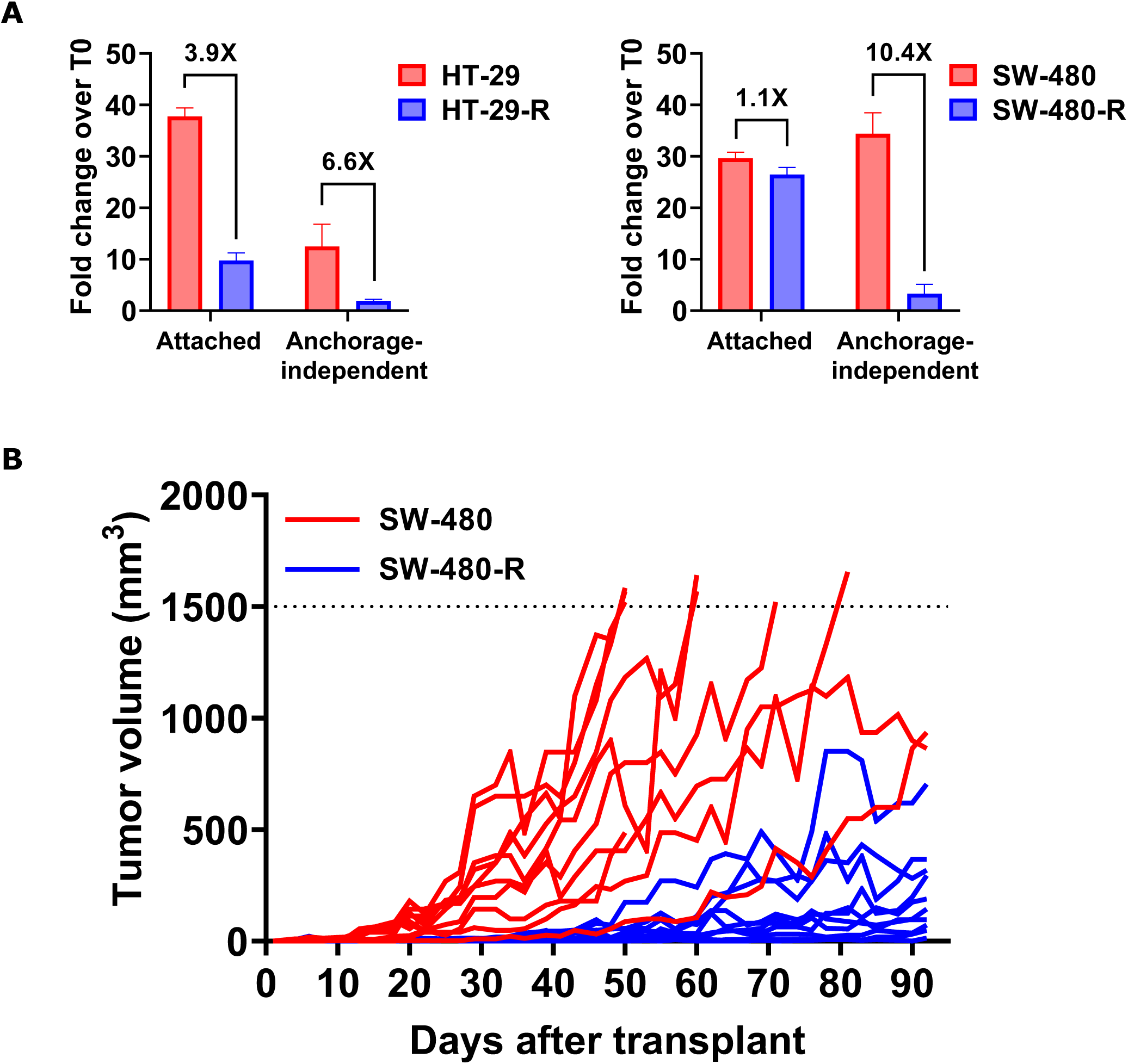
Acquired resistance to the combination of LB-100 and Adavosertib suppress anchorage-independent and tumor growth in CRC models. (A) Endpoint proliferation of HT-29 and SW-480 parental and resistant cells growing attached or in anchorage-independent conditions. Cells were plated in the same density on regular or cell-repellent culture plates in the absence of drugs and grown for 5-6 days. Proliferation was addressed by CellTiter-Glo 3D^®^ and is expressed as fold change over T0. **(B)** Growth curves of the individual tumors from SW-480 parental and resistant cells measured 3 times per week. The dashed line indicates the 1500 mm^3^ ethical sacrifice.

## SUPPLEMENTARY TABLE LEGENDS

**Supplementary table 1:** Cell lines with oncogenic drivers. The mutational status of the cell lines was compiled from the ATCC, Catalogue of Somatic Mutations in Cancer (COSMIC) and Cell Model Passport, Wellcome Trust Sanger Institute, and Depmap portal databases.

| Cancer type | Cell line | Oncogenic drivers | MSI status |
| --- | --- | --- | --- |
| Colorectal | DiFi | EGFR amplification, TP53<sup>K132R</sup>, APC<sup>E1151*</sup> | MSS |
|  | DLD1 | KRAS<sup>G13D</sup>, APC<sup>R2166*</sup>, PIK3CA<sup>E545K/D549N</sup>, TP53<sup>S241F</sup> | MSI |
|  | HCT-116 | KRAS<sup>G13D</sup>, PIK3CA<sup>H1047R</sup>, | MSI |
|  | HT-29 | BRAF<sup>V600E</sup>, TP53<sup>R273H</sup>, APC<sup>E853*</sup>, PIK3CA<sup>P449T</sup>, SMAD4<sup>Q311*</sup> | MSS |
|  | LoVo | KRAS<sup>G13D</sup>, APC<sup>R1114*</sup>, FBXW7<sup>R505C</sup> | MSI |
|  | RKO | BRAF<sup>V600E</sup>, PIK3CA<sup>H1047R</sup> | MSI |
|  | SW-480 | KRAS<sup>G12V</sup>, TP53<sup>R273H/P309S</sup>, APC<sup>Q1338*</sup> | MSS |
| Pancreatic | AspC-1 | KRAS<sup>G12D</sup>, TP53<sup>C135fs*35</sup>, CDKN2A<sup>L78fs*41</sup>, SMAD4<sup>R100T</sup> | MSS |
|  | MIA-PaCa-2 | KRAS<sup>G12C</sup>, TP53<sup>R248W</sup>, CDKN2A<sup>Del</sup> | MSS |
|  | PANC-1 | KRAS<sup>G12D</sup>, TP53<sup>R273H</sup> | MSS |
|  | YAPC | KRAS<sup>G12V</sup>, TP53<sup>H179R</sup>, SMAD4<sup>R515fs*22</sup>, CDKN2A<sup>Del</sup> | Unknown |
| Cholangiocarcinoma | EGI-1 | KRAS<sup>G12D</sup>, TP53<sup>R273H</sup> | Unknown |
|  | HuCC-T1 | KRAS<sup>G12D</sup>, TP53<sup>R175H</sup>, FBXW7<sup>S294*</sup> | MSS |
|  | RBE | KRAS<sup>G12V</sup> | Unknown |
|  | SSP-25 | TP53<sup>R175H</sup> | Unknown |

**Supplementary table 2:**
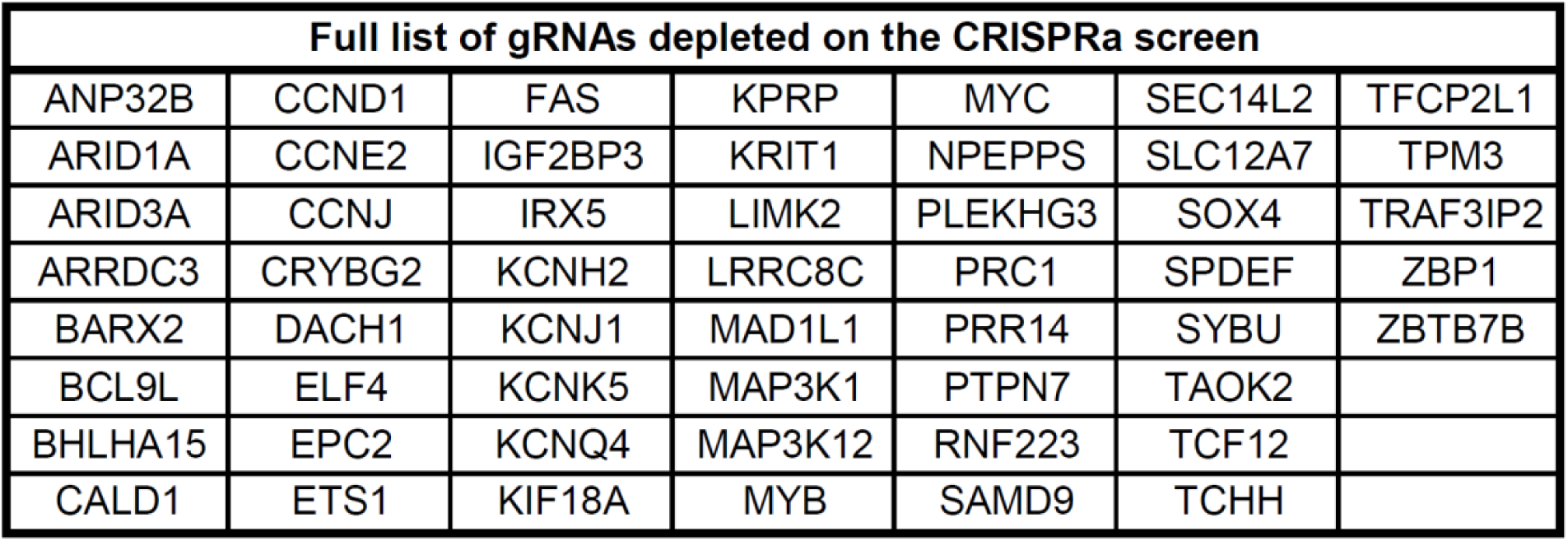
Full list of genes whose overexpression was selectively toxic in the presence of LB-100 in HT-29 cells in the CRISPRa screen. FDR smaller or equal to 0.25 and log2 fold change smaller or equal to -1 in treated/untreated comparison were criteria for hit selection.

| Full list of gRNAs depleted on the CRISPRa screen |  |  |  |  |  |  |
| --- | --- | --- | --- | --- | --- | --- |
| ANP32B | CCND1 | FAS | KPRP | MYC | SEC14L2 | TFCP2L1 |
| ARID1A | CCNE2 | IGF2BP3 | KRIT1 | NPEPPS | SLC12A7 | TPM3 |
| ARID3A | CCNJ | IRX5 | LIMK2 | PLEKHG3 | SOX4 | TRAF3IP2 |
| ARRDC3 | CRYBG2 | KCNH2 | LRRC8C | PRC1 | SPDEF | ZBP1 |
| BARX2 | DACH1 | KCNJ1 | MAD1L1 | PRR14 | SYBU | ZBTB7B |
| BCL9L | ELF4 | KCNK5 | MAP3K1 | PTPN7 | TAOK2 |  |
| BHLHA15 | EPC2 | KCNQ4 | MAP3K12 | RNF223 | TCF12 |  |
| CALD1 | ETS1 | KIF18A | MYB | SAMD9 | TCHH |  |

**Supplementary table 3:** Full list of genes whose knockout attenuated LB-100 toxicity in SW-480 cells in the CRISPR-KO screen. FDR smaller or equal to 0.25 and log2 fold change greater or equal to 1 in treated/untreated comparison were criteria for hit selection.

| Full list of gRNA enriched on the CRISPR-KO screen |  |  |  |  |  |  |
| --- | --- | --- | --- | --- | --- | --- |
| AMOTL2 | DNM2 | GPR137 | MED12 | PLA2G3 | SUV39H1 | ZC3HAV1 |
| ARHGEF15 | EGR4 | LEF1 | MED15 | PROX1 | TADA1 | ZXDA |
| BCL9L | EP300 | MAPK1 | PAWR | RGS7BP | TYW5 |  |
| CTNNB1 | GPI | MAPK14 | PEBP1 | RTFDC1 | UBE2Q1 |  |

**Supplementary table 4:**
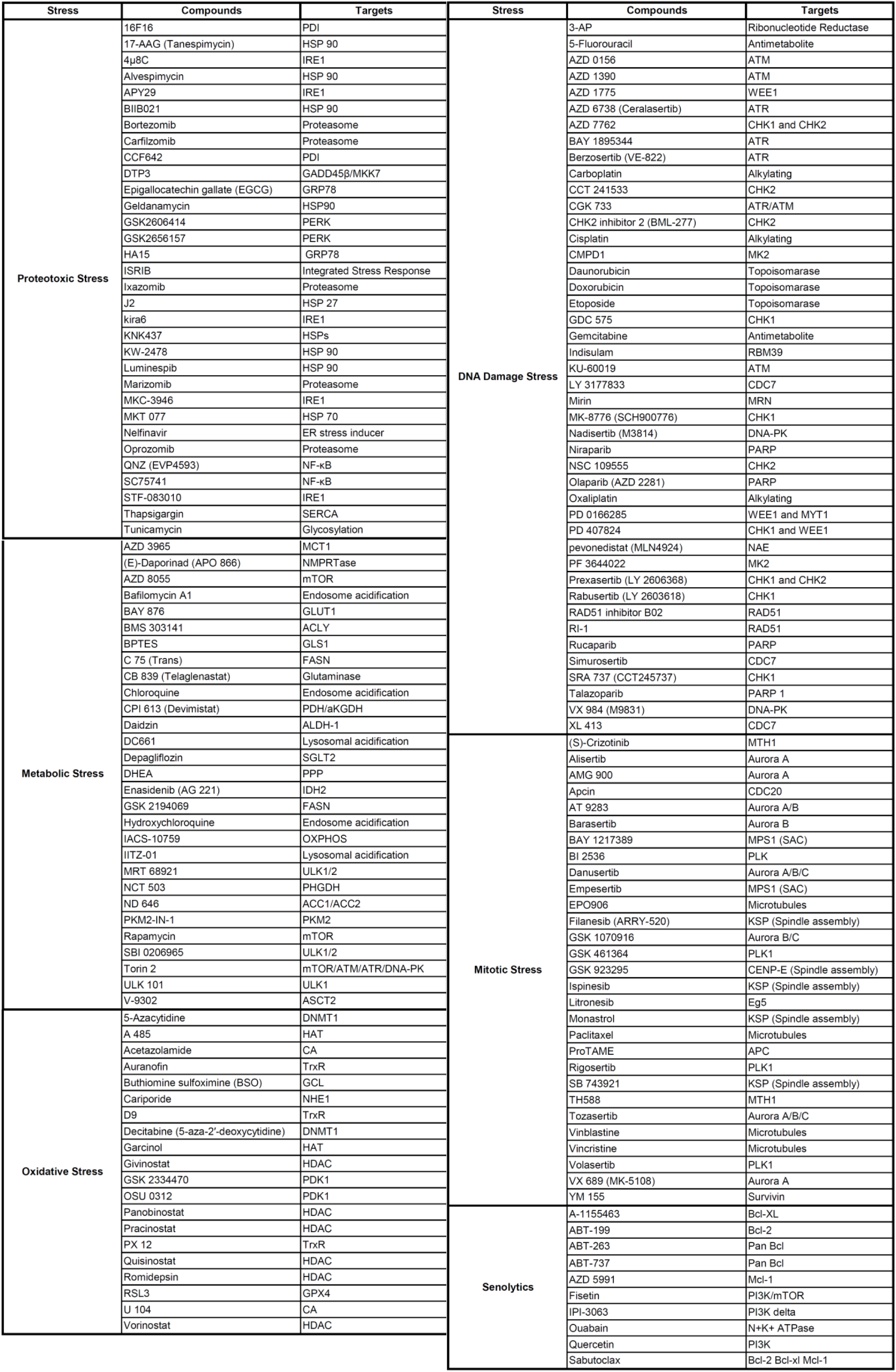
The composition of the stress-focused drug library. Compounds comprising the stress-focused drug library with their respective targets.

**Supplementary table 5:**
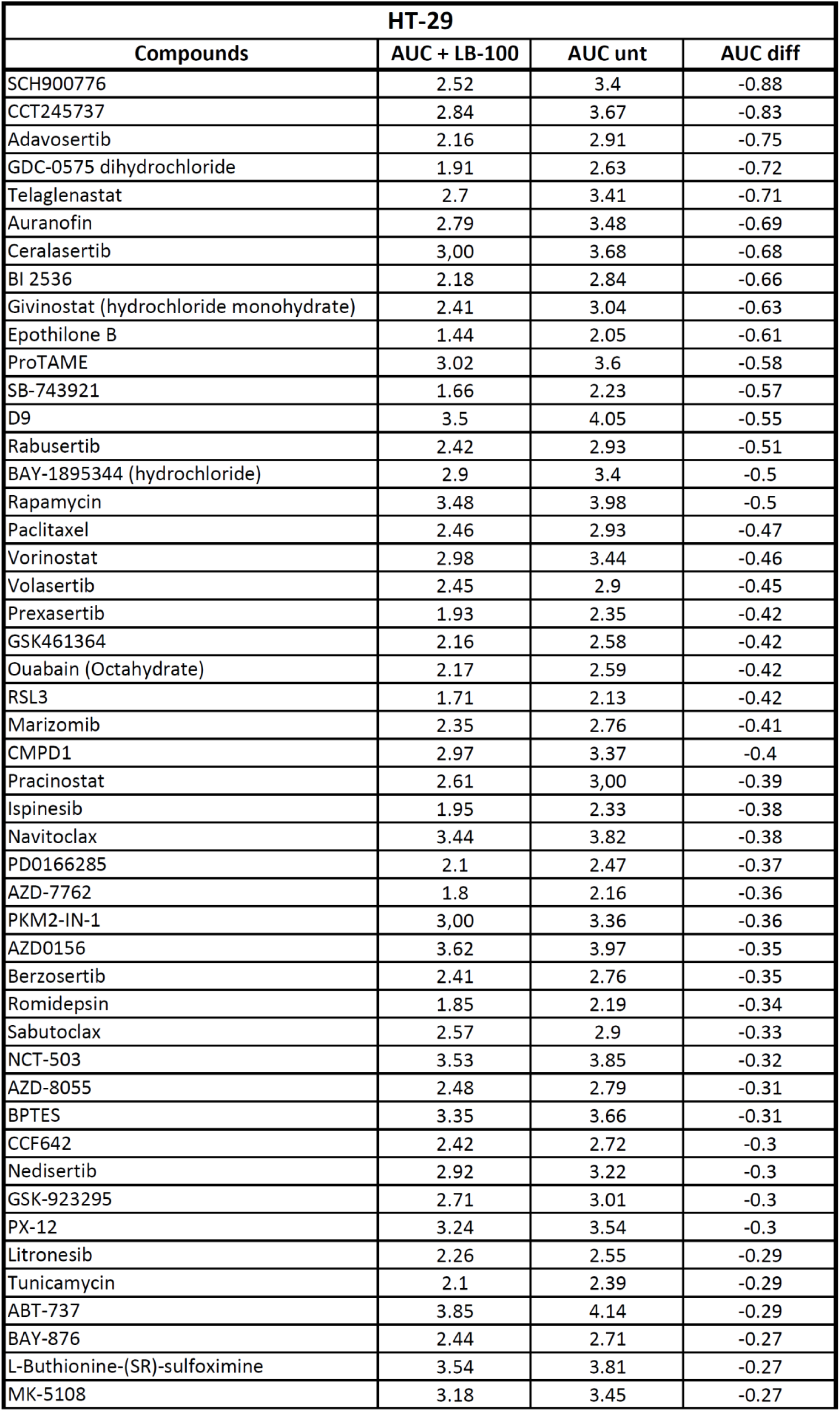
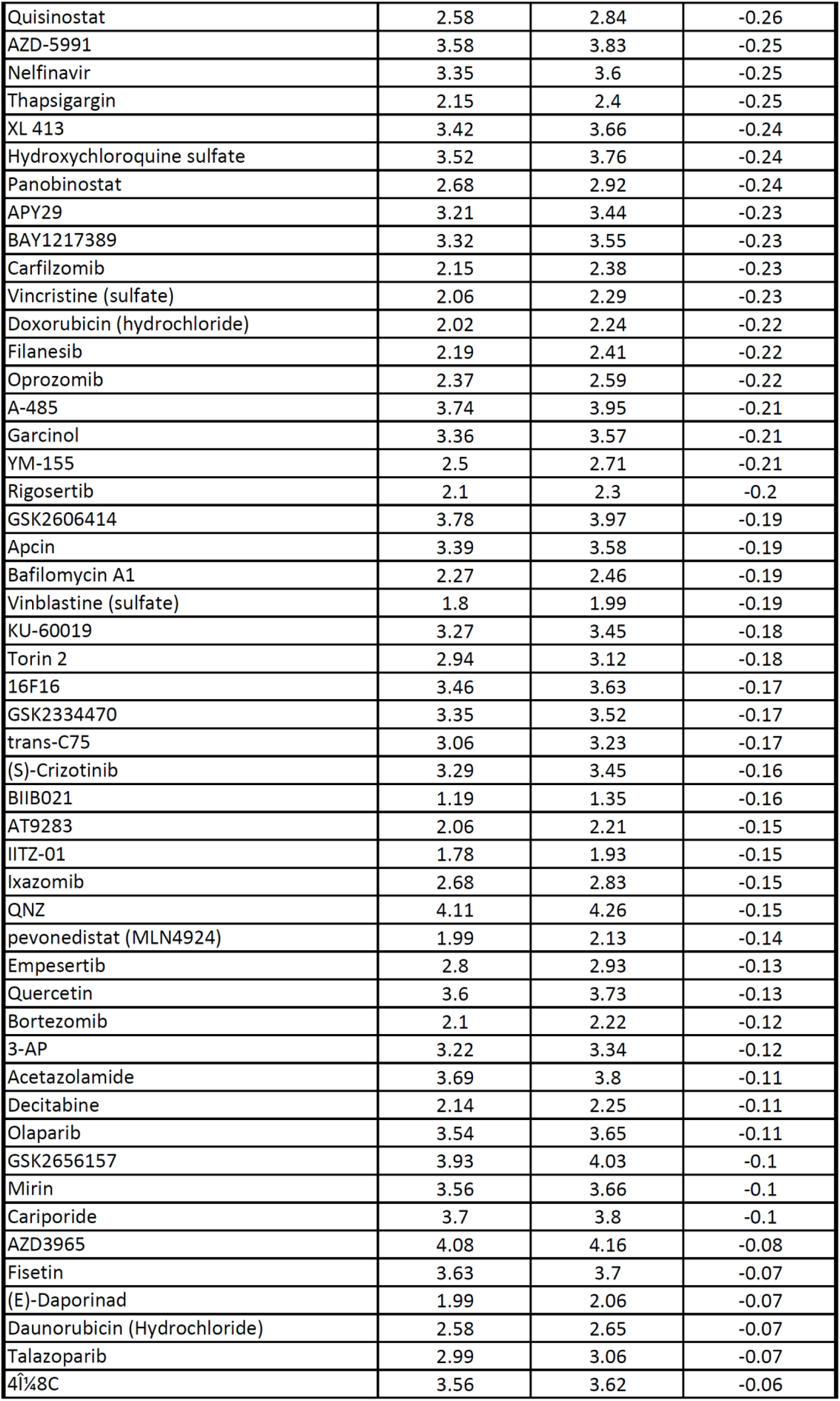
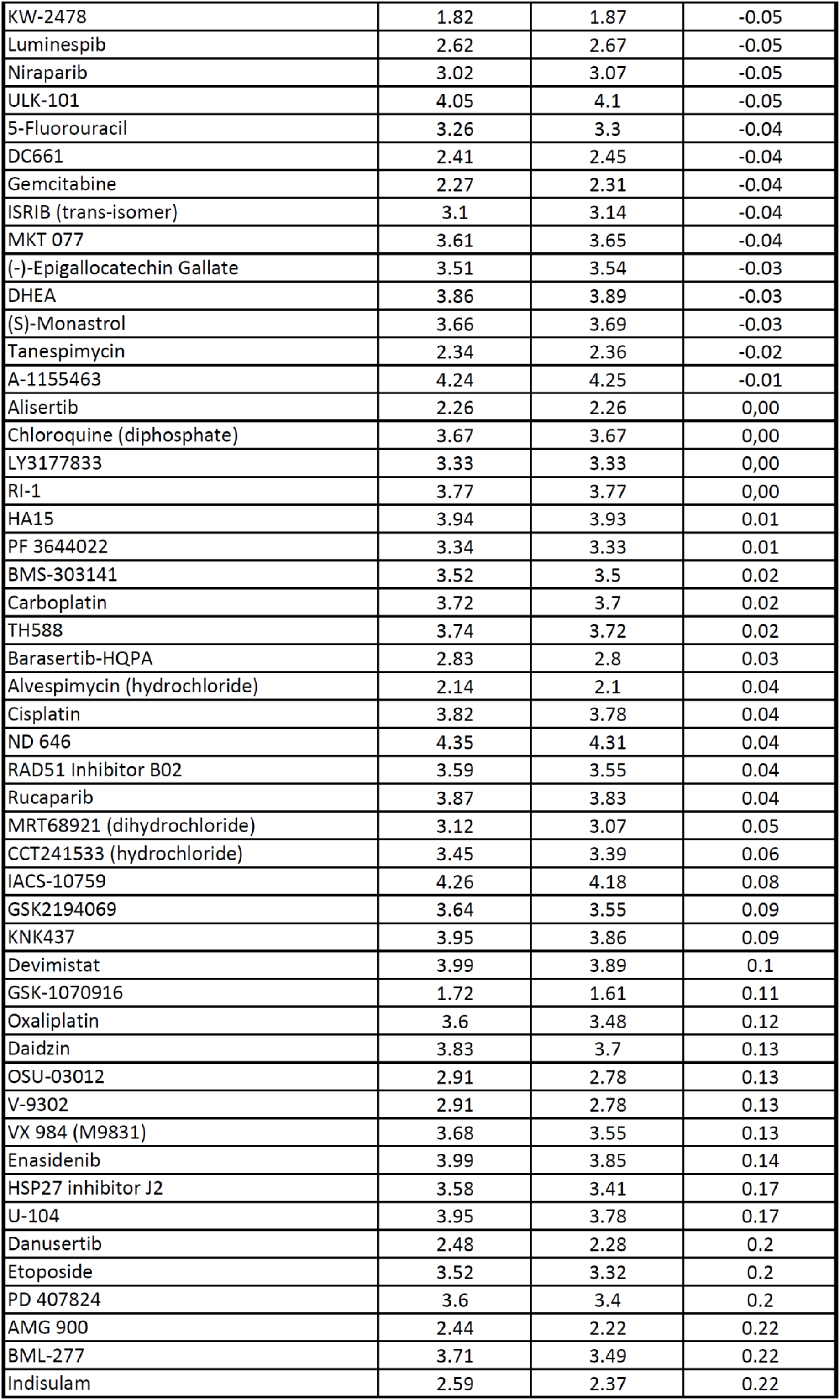
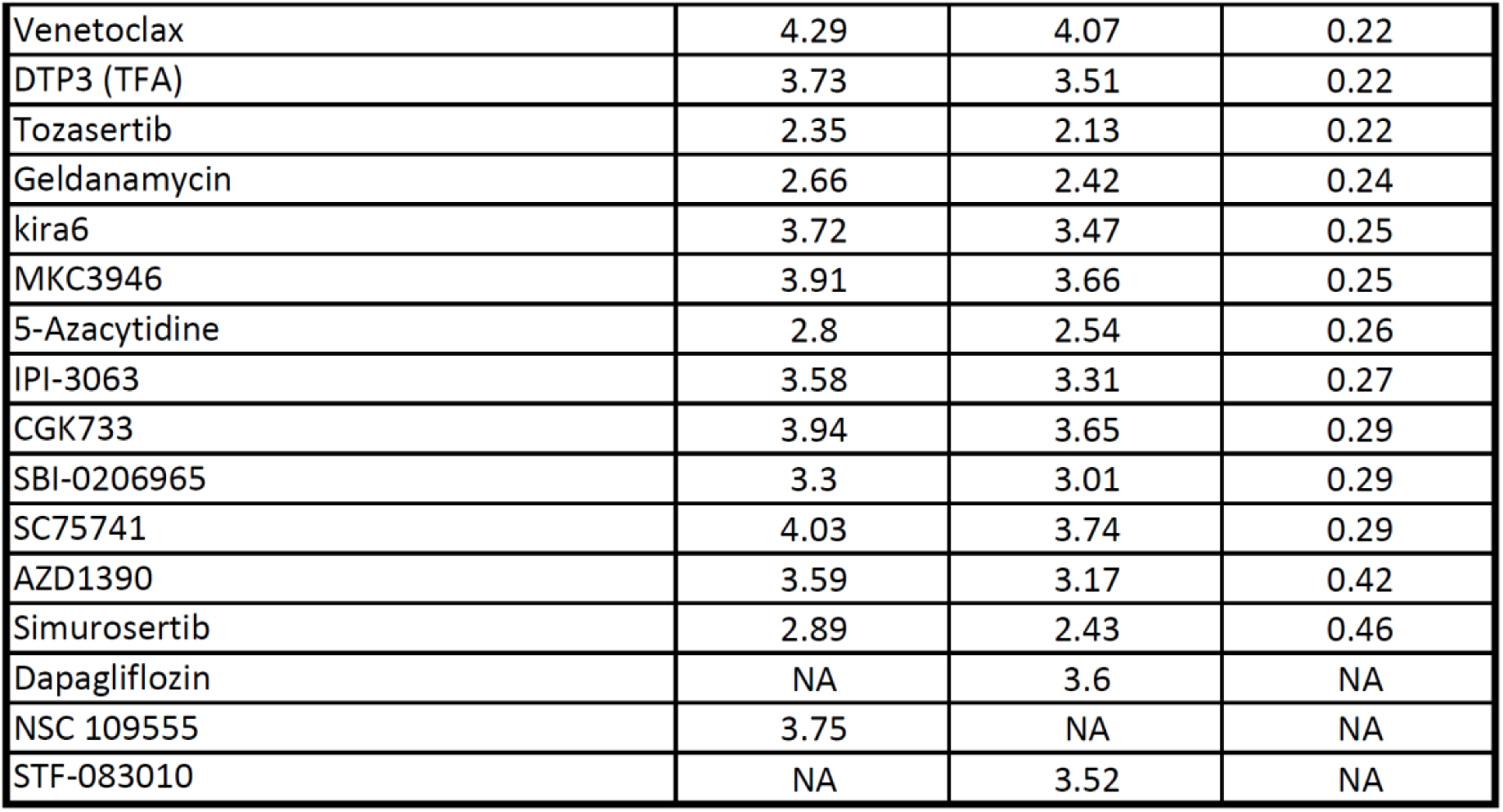
Stress-focused drug screens AUC differences in HT-29 cells. Area Under the Curve (AUC) from each compound of the stress-focused drug screen in the presence or absence of LB-100. Compounds are ranked by the difference in the AUC between LB-100-treated and untreated samples.

| HT-29 |  |  |  |
| --- | --- | --- | --- |
| Compounds | AUC + LB-100 | AUC unt | AUC diff |
| SCH900776 | 2.52 | 3.4 | -0.88 |
| CCT245737 | 2.84 | 3.67 | -0.83 |
| Adavosertib | 2.16 | 2.91 | -0.75 |
| GDC-0575 dihydrochloride | 1.91 | 2.63 | -0.72 |
| Telaglenastat | 2.7 | 3.41 | -0.71 |
| Auranofin | 2.79 | 3.48 | -0.69 |
| Ceralasertib | 3,00 | 3.68 | -0.68 |
| BI 2536 | 2.18 | 2.84 | -0.66 |
| Givinostat (hydrochloride monohydrate) | 2.41 | 3.04 | -0.63 |
| Epothilone B | 1.44 | 2.05 | -0.61 |
| ProTAME | 3.02 | 3.6 | -0.58 |
| SB-743921 | 1.66 | 2.23 | -0.57 |
| D9 | 3.5 | 4.05 | -0.55 |
| Rabusertib | 2.42 | 2.93 | -0.51 |
| BAY-1895344 (hydrochloride) | 2.9 | 3.4 | -0.5 |
| Rapamycin | 3.48 | 3.98 | -0.5 |
| Paclitaxel | 2.46 | 2.93 | -0.47 |
| Vorinostat | 2.98 | 3.44 | -0.46 |
| Volasertib | 2.45 | 2.9 | -0.45 |
| Prexasertib | 1.93 | 2.35 | -0.42 |
| GSK461364 | 2.16 | 2.58 | -0.42 |
| Ouabain (Octahydrate) | 2.17 | 2.59 | -0.42 |
| RSL3 | 1.71 | 2.13 | -0.42 |
| Marizomib | 2.35 | 2.76 | -0.41 |
| CMPD1 | 2.97 | 3.37 | -0.4 |
| Pracinostat | 2.61 | 3,00 | -0.39 |
| Ispinesib | 1.95 | 2.33 | -0.38 |
| Navitoclax | 3.44 | 3.82 | -0.38 |
| PD0166285 | 2.1 | 2.47 | -0.37 |
| AZD-7762 | 1.8 | 2.16 | -0.36 |
| PKM2-IN-1 | 3,00 | 3.36 | -0.36 |
| AZD0156 | 3.62 | 3.97 | -0.35 |
| Berzosertib | 2.41 | 2.76 | -0.35 |
| Romidepsin | 1.85 | 2.19 | -0.34 |
| Sabutoclax | 2.57 | 2.9 | -0.33 |
| NCT-503 | 3.53 | 3.85 | -0.32 |
| AZD-8055 | 2.48 | 2.79 | -0.31 |
| BPTES | 3.35 | 3.66 | -0.31 |
| CCF642 | 2.42 | 2.72 | -0.3 |
| Nedisertib | 2.92 | 3.22 | -0.3 |
| GSK-923295 | 2.71 | 3.01 | -0.3 |
| PX-12 | 3.24 | 3.54 | -0.3 |
| Litronesib | 2.26 | 2.55 | -0.29 |
| Tunicamycin | 2.1 | 2.39 | -0.29 |
| ABT-737 | 3.85 | 4.14 | -0.29 |
| BAY-876 | 2.44 | 2.71 | -0.27 |
| L-Buthionine-(SR)-sulfoximine | 3.54 | 3.81 | -0.27 |
| MK-5108 | 3.18 | 3.45 | -0.27 |

Supplementary table S5
|  |  |  |  |
| --- | --- | --- | --- |
| Quisinostat | 2.58 | 2.84 | -0.26 |
| AZD-5991 | 3.58 | 3.83 | -0.25 |
| Nelfinavir | 3.35 | 3.6 | -0.25 |
| Thapsigargin | 2.15 | 2.4 | -0.25 |
| XL 413 | 3.42 | 3.66 | -0.24 |
| Hydroxychloroquine sulfate | 3.52 | 3.76 | -0.24 |
| Panobinostat | 2.68 | 2.92 | -0.24 |
| APY29 | 3.21 | 3.44 | -0.23 |
| BAY1217389 | 3.32 | 3.55 | -0.23 |
| Carfilzomib | 2.15 | 2.38 | -0.23 |
| Vincristine (sulfate) | 2.06 | 2.29 | -0.23 |
| Doxorubicin (hydrochloride) | 2.02 | 2.24 | -0.22 |
| Filanesib | 2.19 | 2.41 | -0.22 |
| Oprozomib | 2.37 | 2.59 | -0.22 |
| A-485 | 3.74 | 3.95 | -0.21 |
| Garcinol | 3.36 | 3.57 | -0.21 |
| YM-155 | 2.5 | 2.71 | -0.21 |
| Rigosertib | 2.1 | 2.3 | -0.2 |
| GSK2606414 | 3.78 | 3.97 | -0.19 |
| Apcin | 3.39 | 3.58 | -0.19 |
| Bafilomycin A1 | 2.27 | 2.46 | -0.19 |
| Vinblastine (sulfate) | 1.8 | 1.99 | -0.19 |
| KU-60019 | 3.27 | 3.45 | -0.18 |
| Torin 2 | 2.94 | 3.12 | -0.18 |
| 16F16 | 3.46 | 3.63 | -0.17 |
| GSK2334470 | 3.35 | 3.52 | -0.17 |
| trans-C75 | 3.06 | 3.23 | -0.17 |
| (S)-Crizotinib | 3.29 | 3.45 | -0.16 |
| BIIB021 | 1.19 | 1.35 | -0.16 |
| AT9283 | 2.06 | 2.21 | -0.15 |
| IITZ-01 | 1.78 | 1.93 | -0.15 |
| Ixazomib | 2.68 | 2.83 | -0.15 |
| QNZ | 4.11 | 4.26 | -0.15 |
| pevonedistat (MLN4924) | 1.99 | 2.13 | -0.14 |
| Empesertib | 2.8 | 2.93 | -0.13 |
| Quercetin | 3.6 | 3.73 | -0.13 |
| Bortezomib | 2.1 | 2.22 | -0.12 |
| 3-AP | 3.22 | 3.34 | -0.12 |
| Acetazolamide | 3.69 | 3.8 | -0.11 |
| Decitabine | 2.14 | 2.25 | -0.11 |
| Olaparib | 3.54 | 3.65 | -0.11 |
| GSK2656157 | 3.93 | 4.03 | -0.1 |
| Mirin | 3.56 | 3.66 | -0.1 |
| Cariporide | 3.7 | 3.8 | -0.1 |
| AZD3965 | 4.08 | 4.16 | -0.08 |
| Fisetin | 3.63 | 3.7 | -0.07 |
| (E)-Daporinad | 1.99 | 2.06 | -0.07 |
| Daunorubicin (Hydrochloride) | 2.58 | 2.65 | -0.07 |
| Talazoparib | 2.99 | 3.06 | -0.07 |
| 4Î¼8C | 3.56 | 3.62 | -0.06 |

Supplementary table S5
|  |  |  |  |
| --- | --- | --- | --- |
| KW-2478 | 1.82 | 1.87 | -0.05 |
| Luminespib | 2.62 | 2.67 | -0.05 |
| Niraparib | 3.02 | 3.07 | -0.05 |
| ULK-101 | 4.05 | 4.1 | -0.05 |
| 5-Fluorouracil | 3.26 | 3.3 | -0.04 |
| DC661 | 2.41 | 2.45 | -0.04 |
| Gemcitabine | 2.27 | 2.31 | -0.04 |
| ISRIB (trans-isomer) | 3.1 | 3.14 | -0.04 |
| MKT 077 | 3.61 | 3.65 | -0.04 |
| (-)-Epigallocatechin Gallate | 3.51 | 3.54 | -0.03 |
| DHEA | 3.86 | 3.89 | -0.03 |
| (S)-Monastrol | 3.66 | 3.69 | -0.03 |
| Tanespimycin | 2.34 | 2.36 | -0.02 |
| A-1155463 | 4.24 | 4.25 | -0.01 |
| Alisertib | 2.26 | 2.26 | 0,00 |
| Chloroquine (diphosphate) | 3.67 | 3.67 | 0,00 |
| LY3177833 | 3.33 | 3.33 | 0,00 |
| RI-1 | 3.77 | 3.77 | 0,00 |
| HA15 | 3.94 | 3.93 | 0.01 |
| PF 3644022 | 3.34 | 3.33 | 0.01 |
| BMS-303141 | 3.52 | 3.5 | 0.02 |
| Carboplatin | 3.72 | 3.7 | 0.02 |
| TH588 | 3.74 | 3.72 | 0.02 |
| Barasertib-HQPA | 2.83 | 2.8 | 0.03 |
| Alvespimycin (hydrochloride) | 2.14 | 2.1 | 0.04 |
| Cisplatin | 3.82 | 3.78 | 0.04 |
| ND 646 | 4.35 | 4.31 | 0.04 |
| RAD51 Inhibitor B02 | 3.59 | 3.55 | 0.04 |
| Rucaparib | 3.87 | 3.83 | 0.04 |
| MRT68921 (dihydrochloride) | 3.12 | 3.07 | 0.05 |
| CCT241533 (hydrochloride) | 3.45 | 3.39 | 0.06 |
| IACS-10759 | 4.26 | 4.18 | 0.08 |
| GSK2194069 | 3.64 | 3.55 | 0.09 |
| KNK437 | 3.95 | 3.86 | 0.09 |
| Devimistat | 3.99 | 3.89 | 0.1 |
| GSK-1070916 | 1.72 | 1.61 | 0.11 |
| Oxaliplatin | 3.6 | 3.48 | 0.12 |
| Daidzin | 3.83 | 3.7 | 0.13 |
| OSU-03012 | 2.91 | 2.78 | 0.13 |
| V-9302 | 2.91 | 2.78 | 0.13 |
| VX 984 (M9831) | 3.68 | 3.55 | 0.13 |
| Enasidenib | 3.99 | 3.85 | 0.14 |
| HSP27 inhibitor J2 | 3.58 | 3.41 | 0.17 |
| U-104 | 3.95 | 3.78 | 0.17 |
| Danuserib | 2.48 | 2.28 | 0.2 |
| Etoposide | 3.52 | 3.32 | 0.2 |
| PD 407824 | 3.6 | 3.4 | 0.2 |
| AMG 900 | 2.44 | 2.22 | 0.22 |
| BML-277 | 3.71 | 3.49 | 0.22 |
| Indisulam | 2.59 | 2.37 | 0.22 |

Supplementary table S5
|  |  |  |  |
| --- | --- | --- | --- |
| Venetoclax | 4.29 | 4.07 | 0.22 |
| DTP3 (TFA) | 3.73 | 3.51 | 0.22 |
| Tozasertib | 2.35 | 2.13 | 0.22 |
| Geldanamycin | 2.66 | 2.42 | 0.24 |
| kira6 | 3.72 | 3.47 | 0.25 |
| MKC3946 | 3.91 | 3.66 | 0.25 |
| 5-Azacytidine | 2.8 | 2.54 | 0.26 |
| IPI-3063 | 3.58 | 3.31 | 0.27 |
| CGK733 | 3.94 | 3.65 | 0.29 |
| SBI-0206965 | 3.3 | 3.01 | 0.29 |
| SC75741 | 4.03 | 3.74 | 0.29 |
| AZD1390 | 3.59 | 3.17 | 0.42 |
| Simurosertib | 2.89 | 2.43 | 0.46 |
| Dapagliflozin | NA | 3.6 | NA |
| NSC 109555 | 3.75 | NA | NA |
| STF-083010 | NA | 3.52 | NA |

**Supplementary table 6:** Stress-focused drug screens AUC differences in SW-480 cells. Area Under the Curve (AUC) from each compound of the stress-focused drug screen in the presence or absence of LB-100. Compounds are ranked by the difference in the AUC between LB-100-treated and untreated samples.

| SW-480 |  |  |  |
| --- | --- | --- | --- |
| Compounds | AUC + LB-100 | AUC unt | AUC diff |
| A-1155463 | 1.65 | 2.77 | -1.12 |
| Adavosertib | 2.22 | 2.8 | -0.58 |
| Rabusertib | 2.66 | 3.14 | -0.48 |
| Navitoclax | 1.88 | 2.32 | -0.44 |
| BI 2536 | 2.29 | 2.69 | -0.4 |
| PD0166285 | 1.83 | 2.23 | -0.4 |
| GDC-0575 dihydrochloride | 2.35 | 2.72 | -0.37 |
| Prexasertib | 2.29 | 2.65 | -0.36 |
| Telaglenastat | 3.22 | 3.51 | -0.29 |
| CCT245737 | 2.98 | 3.25 | -0.27 |
| Marizomib | 2,00 | 2.25 | -0.25 |
| Rigosertib | 1.97 | 2.21 | -0.24 |
| BPTES | 3.17 | 3.41 | -0.24 |
| GSK461364 | 2.32 | 2.53 | -0.21 |
| RSL3 | 2.51 | 2.71 | -0.2 |
| Vincristine (sulfate) | 2.2 | 2.4 | -0.2 |
| Volasertib | 2.8 | 2.99 | -0.19 |
| SCH900776 | 2.97 | 3.16 | -0.19 |
| Carfilzomib | 1.85 | 2.03 | -0.18 |
| Pracinostat | 2.04 | 2.22 | -0.18 |
| Carboplatin | 3.12 | 3.28 | -0.16 |
| Sabutoclax | 2.49 | 2.65 | -0.16 |
| CCF642 | 1.7 | 1.85 | -0.15 |
| Rapamycin | 2.67 | 2.81 | -0.14 |
| Givinostat (hydrochloride monohydrate) | 1.96 | 2.08 | -0.12 |
| Alvespimycin (hydrochloride) | 2.62 | 2.72 | -0.1 |
| D9 | 2.44 | 2.54 | -0.1 |
| Oprozomib | 2.27 | 2.37 | -0.1 |
| YM-155 | 1.81 | 1.9 | -0.09 |
| Bortezomib | 1.52 | 1.61 | -0.09 |
| Torin 2 | 2.36 | 2.45 | -0.09 |
| AZD-8055 | 1.97 | 2.06 | -0.09 |
| Nelfinavir | 3.39 | 3.47 | -0.08 |
| U-104 | 3.14 | 3.22 | -0.08 |
| Luminespib | 2.26 | 2.33 | -0.07 |
| Panobinostat | 2.07 | 2.13 | -0.06 |
| 4Î¼8C | 3.47 | 3.53 | -0.06 |
| BAY-876 | 3.52 | 3.57 | -0.05 |
| Ceralasertib | 2.98 | 3.03 | -0.05 |
| Ouabain (Octahydrate) | 2.17 | 2.22 | -0.05 |
| (-)-Epigallocatechin Gallate | 2.57 | 2.61 | -0.04 |
| PD 407824 | 3.2 | 3.23 | -0.03 |
| MKC3946 | 3.29 | 3.32 | -0.03 |
| Dapagliflozin | 3.44 | 3.47 | -0.03 |
| IITZ-01 | 2.37 | 2.39 | -0.02 |
| Epothilone B | 1.92 | 1.94 | -0.02 |
| 16F16 | 3.16 | 3.18 | -0.02 |
| Talazoparib | 3.25 | 3.26 | -0.01 |

Supplementary table S6
|  |  |  |  |
| --- | --- | --- | --- |
| Ixazomib | 2.29 | 2.3 | -0.01 |
| A-485 | 3.04 | 3.05 | -0.01 |
| Vinblastine (sulfate) | 2.27 | 2.27 | 0,00 |
| Thapsigargin | 1.94 | 1.94 | 0,00 |
| Garcinol | 2.84 | 2.84 | 0,00 |
| Auranofin | 1.9 | 1.9 | 0,00 |
| Chloroquine (diphosphate) | 3.21 | 3.21 | 0,00 |
| PKM2-IN-1 | 2.52 | 2.52 | 0,00 |
| GSK2656157 | 3.13 | 3.13 | 0,00 |
| BAY-1895344 (hydrochloride) | 3.01 | 3,00 | 0.01 |
| HSP27 inhibitor J2 | 3.35 | 3.34 | 0.01 |
| RAD51 Inhibitor B02 | 3.52 | 3.51 | 0.01 |
| OSU-03012 | 2.82 | 2.81 | 0.01 |
| Vorinostat | 2.51 | 2.5 | 0.01 |
| Tanespimycin | 2.95 | 2.92 | 0.03 |
| Niraparib | 2.78 | 2.75 | 0.03 |
| GSK-923295 | 3.01 | 2.98 | 0.03 |
| AZD0156 | 3.34 | 3.31 | 0.03 |
| Quisinostat | 2.09 | 2.05 | 0.04 |
| IACS-10759 | 3.32 | 3.28 | 0.04 |
| V-9302 | 2.82 | 2.78 | 0.04 |
| KU-60019 | 3.42 | 3.38 | 0.04 |
| SBI-0206965 | 3.51 | 3.46 | 0.05 |
| Cisplatin | 3,00 | 2.95 | 0.05 |
| Bafilomycin A1 | 1.34 | 1.29 | 0.05 |
| AZD-7762 | 2.08 | 2.02 | 0.06 |
| QNZ | 3.06 | 3,00 | 0.06 |
| (E)-Daporinad | 2.19 | 2.13 | 0.06 |
| Romidepsin | 1.22 | 1.15 | 0.07 |
| Berzosertib | 2.95 | 2.88 | 0.07 |
| GSK2334470 | 3.3 | 3.23 | 0.07 |
| L-Buthionine-(SR)-sulfoximine | 3.4 | 3.32 | 0.08 |
| SB-743921 | 2.47 | 2.39 | 0.08 |
| XL 413 | 3.54 | 3.46 | 0.08 |
| Rucaparib | 3.27 | 3.19 | 0.08 |
| AZD-5991 | 3.38 | 3.3 | 0.08 |
| KW-2478 | 2.95 | 2.86 | 0.09 |
| Ispinesib | 2.67 | 2.57 | 0.1 |
| Filanesib | 2.66 | 2.56 | 0.1 |
| Barasertib-HQPA | 3.6 | 3.5 | 0.1 |
| BIIB021 | 2.03 | 1.93 | 0.1 |
| IPI-3063 | 3.66 | 3.56 | 0.1 |
| kira6 | 3.47 | 3.37 | 0.1 |
| GSK2606414 | 3.39 | 3.29 | 0.1 |
| PX-12 | 3.21 | 3.11 | 0.1 |
| Daidzin | 3.35 | 3.24 | 0.11 |
| Gemcitabine | 2.38 | 2.27 | 0.11 |
| HA15 | 3.47 | 3.36 | 0.11 |
| Quercetin | 3.25 | 3.13 | 0.12 |
| AZD3965 | 3.42 | 3.3 | 0.12 |

Supplementary table S6
|  |  |  |  |
| --- | --- | --- | --- |
| DC661 | 2.33 | 2.21 | 0.12 |
| (S)-Crizotinib | 3.22 | 3.09 | 0.13 |
| TH588 | 3.68 | 3.54 | 0.14 |
| Paclitaxel | 2.83 | 2.68 | 0.15 |
| BMS-303141 | 3.15 | 3.00 | 0.15 |
| trans-C75 | 2.82 | 2.67 | 0.15 |
| Devimistat | 3.29 | 3.14 | 0.15 |
| 3-AP | 2.99 | 2.84 | 0.15 |
| Daunorubicin (Hydrochloride) | 2.56 | 2.4 | 0.16 |
| Fisetin | 3.35 | 3.18 | 0.17 |
| (S)-Monastrol | 3.27 | 3.1 | 0.17 |
| Apcin | 3.44 | 3.27 | 0.17 |
| Doxorubicin (hydrochloride) | 2.09 | 1.92 | 0.17 |
| DTP3 (TFA) | 3.59 | 3.42 | 0.17 |
| CGK733 | 3.51 | 3.33 | 0.18 |
| Venetoclax | 3.53 | 3.35 | 0.18 |
| Acetazolamide | 3.44 | 3.25 | 0.19 |
| Indisulam | 2.99 | 2.8 | 0.19 |
| CCT241533 (hydrochloride) | 3.22 | 3.02 | 0.2 |
| Litronesib | 2.76 | 2.55 | 0.21 |
| 5-Azacytidine | 3.16 | 2.95 | 0.21 |
| 5-Fluorouracil | 3.41 | 3.2 | 0.21 |
| ISRIB (trans-isomer) | 3.65 | 3.44 | 0.21 |
| MKT 077 | 3.3 | 3.09 | 0.21 |
| STF-083010 | 3.32 | 3.1 | 0.22 |
| Nedisertib | 3.28 | 3.06 | 0.22 |
| VX 984 (M9831) | 3.64 | 3.41 | 0.23 |
| NSC 109555 | 3.58 | 3.35 | 0.23 |
| CMPD1 | 2.99 | 2.75 | 0.24 |
| Enasidenib | 3.59 | 3.35 | 0.24 |
| Geldanamycin | 2.61 | 2.37 | 0.24 |
| SC75741 | 3.55 | 3.3 | 0.25 |
| BML-277 | 3.61 | 3.35 | 0.26 |
| Mirin | 2.79 | 2.53 | 0.26 |
| Decitabine | 2.97 | 2.69 | 0.28 |
| Olaparib | 3.58 | 3.3 | 0.28 |
| Oxaliplatin | 2.58 | 2.3 | 0.28 |
| AZD1390 | 3.58 | 3.29 | 0.29 |
| Hydroxychloroquine sulfate | 3.47 | 3.18 | 0.29 |
| Tunicamycin | 2.09 | 1.8 | 0.29 |
| Simurosertib | 3.26 | 2.94 | 0.32 |
| DHEA | 3.38 | 3.06 | 0.32 |
| PF 3644022 | 3.07 | 2.74 | 0.33 |
| ABT-737 | 3.27 | 2.93 | 0.34 |
| Danuserib | 3.15 | 2.81 | 0.34 |
| ULK-101 | 3.49 | 3.14 | 0.35 |
| RI-1 | 3.47 | 3.1 | 0.37 |
| ND 646 | 3.75 | 3.38 | 0.37 |
| Cariporide | 3.31 | 2.94 | 0.37 |
| GSK-1070916 | 3.44 | 3.02 | 0.42 |

Supplementary table S6
|  |  |  |  |
| --- | --- | --- | --- |
| AMG 900 | 3.71 | 3.29 | 0.42 |
| AT9283 | 2.97 | 2.54 | 0.43 |
| MK-5108 | 3.35 | 2.9 | 0.45 |
| Etoposide | 3.29 | 2.82 | 0.47 |
| Tozasertib | 3.59 | 3.11 | 0.48 |
| APY29 | 2.39 | 1.91 | 0.48 |
| LY3177833 | 3.32 | 2.82 | 0.5 |
| BAY1217389 | 3.68 | 3.12 | 0.56 |
| pevonedistat (MLN4924) | 3.42 | 2.85 | 0.57 |
| Empesertib | 3.41 | 2.82 | 0.59 |
| Alisertib | 3.1 | 2.38 | 0.72 |
| MRT68921 (dihydrochloride) | 3.64 | 2.83 | 0.81 |
| ProTAME | NA | 3.38 | NA |
| NCT-503 | NA | 3.29 | NA |
| GSK2194069 | 3.44 | NA | NA |
| KNK437 | 3.5 | NA | NA |

**Supplementary table 7:** Full list of genes whose knockout was selectively toxic in the presence of LB-100 in SW-480 cells in the CRISPR-KO screen. FDR smaller or equal to 0.25 and log2 fold change smaller or equal to -1 in treated/untreated comparison were criteria for hit selection.

| Full list of gRNAs depleted on the CRISPR-KO screen |  |  |  |  |  |
| --- | --- | --- | --- | --- | --- |
| ATPAF2 | GFM2 | MRPL32 | NDUFB4 | PPP1R7 | SRF |
| CYCS | HTT | MRPL41 | PET117 | RAD9A | WEE1 |
| DNAJA3 | KRT18 | MRPS35 | PPP1CA | SFSWAP |  |

